# Truncating *ASXL1* variants rewire cellular metabolism via mitochondrial pyruvate carrier repression

**DOI:** 10.64898/2026.07.13.737346

**Authors:** Isabella Lin, Michael Sigfrid S. Reyes, Abigail S. Krall, Neerja Vashist, Suzanne Sarkissian, Nedas Matulionis, Aileen Ning, Linsey Stiles, Bianca E. Russell, Paul Mark B. Medina, Heather R. Christofk, Valerie A. Arboleda

## Abstract

Bohring-Opitz syndrome (BOS, OMIM#605309) is a rare neurodevelopmental disorder caused by heterozygous and truncating variants in *ASXL1 (Additional Sex Combs Like 1*), a chromatin-associated epigenetic regulator that forms the catalytic PR-DUB complex with BAP1. Truncating *ASXL1* variants are also recurrent somatic drivers in myeloid leukemia, yet the metabolic consequences of these mutations remain undefined. Using patient derived dermal fibroblasts, we show that truncating *ASXL1* variants drive a Warburg-like metabolic state characterized by increased glycolytic flux, and accumulation of pyruvate and lactate. Truncated ASXL1 and BAP1 show aberrant co-occupancy at an H3K4me3-marked intronic regulatory element within *MPC2* intron 1, with broadened ASXL1 occupancy extending beyond BRD4-defined regulatory boundaries while BRD4 positioning remains unchanged, consistent with aberrant PR-DUB complex spreading beyond its normally constrained chromatin territory. This altered occupancy is accompanied by modest but significant reduction in *MPC2* transcript abundance and a disproportionately larger reduction in MPC1 and MPC2 protein levels, indicating that transcriptional dysregulation at this intronic element is amplified at the protein level through post-transcriptional mechanisms including impaired MPC1/MPC2 heterodimer stability. Pharmacologic MPC inhibition recapitulates both the metabolic and Wnt signaling phenotypes of BOS cells, while canonical Wnt activation increases glycolytic flux without reducing MPC abundance, establishing mitochondrial pyruvate restriction as causally upstream of signaling dysregulation. These findings define a previously unrecognized chromatin-to-metabolism axis connecting gain-of-function *ASXL1* truncation to mitochondrial pyruvate transport, identifying MPC as a central mediator of epigenetic-metabolic crosstalk in both a rare developmental syndrome and *ASXL1*-mutant myeloid malignancy.

**Graphical Abstract:** 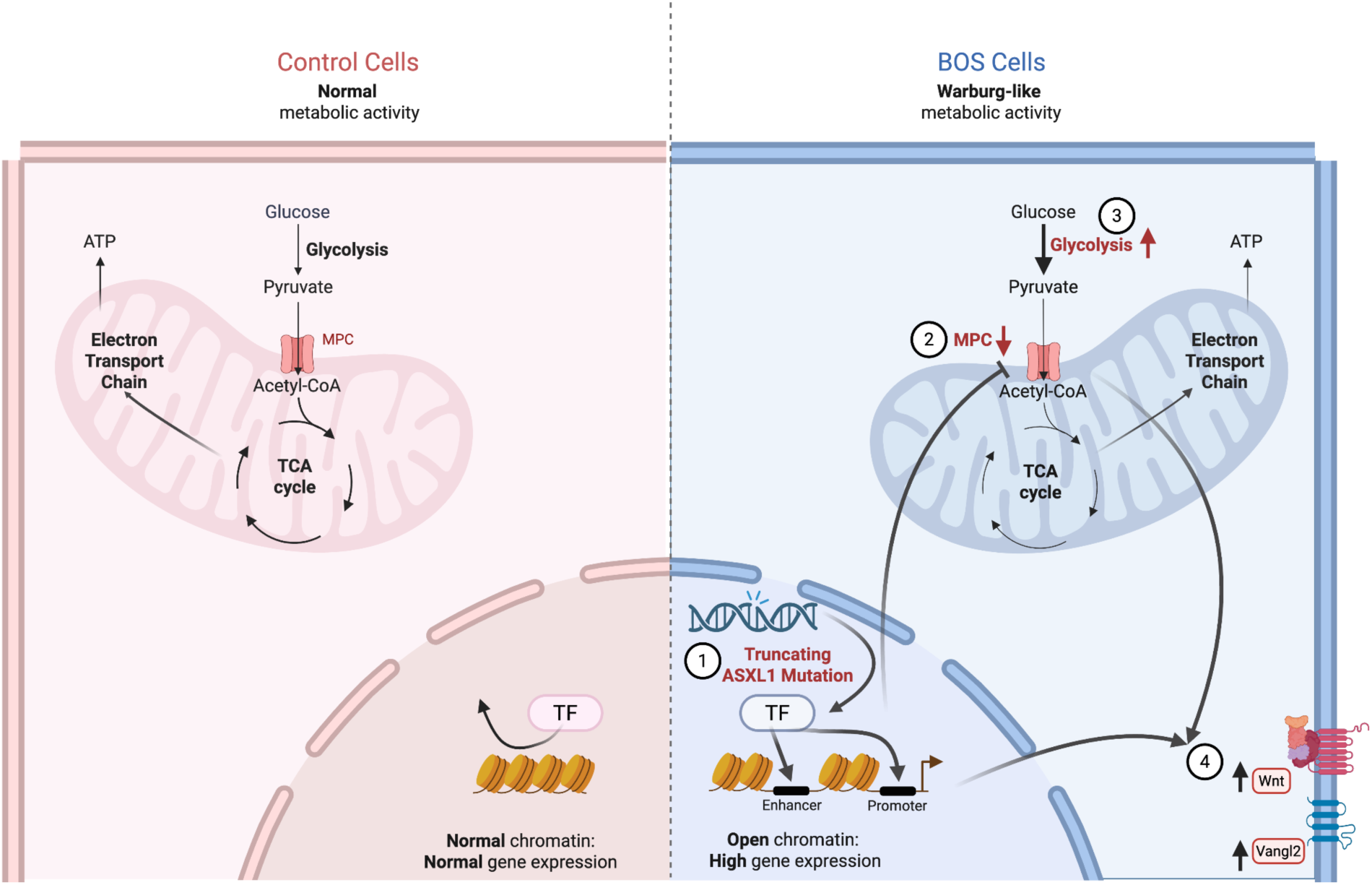

Truncating and heterozygous ASXL1 variants cause a neurodevelopmental syndrome called Bohring-Opitz syndrome. (1) At an epigenetic level, we have shown that Truncating ASXL1 variants drive more open chromatin and aberrant activation of key developmental pathways. (2) Truncating ASXL1 mutations are sufficient to drive Decreased MPC1 and MPC2 protein levels. (3) Decreased MPC1 or MPC2 level or function are sufficient to drive increased glycolysis which is observed in BOS cells. (4) Truncating ASXL1 mutations drive Increased Wnt signaling via MPC depletion. * Increased Wnt signaling (4) is also sufficient to drive increased glycolysis (3), however Increased Wnt signaling does not drive Decreased MPC1 and MPC2 levels (2).

## INTRODUCTION

Bohring-Opitz syndrome (BOS, OMIM #605309) is a rare and severe congenital disorder characterized by profound developmental delays, distinct craniofacial anomalies, growth retardation, and a constellation of clinical manifestations including thick, fast-growing hair, seizures, and recurrent respiratory infections ^1–3^. First described in 1999 by Bohring et al., nearly all molecularly confirmed cases of BOS arise from *de novo* heterozygous truncating germline variants in the *Additional Sex Combs-like 1* (*ASXL1*), a gene known for its role in chromatin regulation, transcriptional control, and development. *ASXL1* is also a frequently observed somatic mutation in myeloid malignancies and independently confers a worse prognosis ^4^. Despite the high frequency of truncating *ASXL1* variants in both BOS and myeloid leukemia, no targeted therapies exist for *ASXL1* variants or its downstream targets.

*ASXL1* encodes a chromatin-associated epigenetic modifier that functions as a scaffold within polycomb repressive complexes (PRCs), which are essential for maintaining cell-type-specific gene expression programs during development ^5–7^. Together with its binding partner BAP1, ASXL1 forms the Polycomb repressive deubiquitinase (PR-DUB) complex, which removes monoubiquitin from H2A lysine 119 (H2AK119ub1) to modulate transcription at Polycomb-regulated loci ^8^. Truncating variants in *ASXL1* generate stable protein fragments that aberrantly enhance BAP1 deubiquitinase activity, producing a hyperactive PR-DUB complex with gain-of-function effects on chromatin state and gene expression ^9,10^. Prior work from our group has demonstrated that ASXL1 truncation drives strong activation of canonical and noncanonical Wnt signaling in both BOS patient-derived fibroblasts and ASXL1-mutant acute myeloid leukemia ^5,11^. Whether these signaling changes are accompanied by, or secondary to, metabolic reprogramming has not been investigated.

Under physiological oxygen conditions, pyruvate derived from glycolysis enters the mitochondrial matrix via the mitochondrial pyruvate carrier (MPC), where it fuels the TCA cycle and oxidative phosphorylation (OXPHOS) for efficient ATP synthesis. Warburg first observed that cancer cells deviate from this program by preferentially using glycolysis even in the presence of oxygen, a phenomenon now termed aerobic glycolysis or the Warburg effect ^12^. This metabolic state is a well-established hallmark of cancer and is increasingly recognized in genetic disorders of epigenetic origin ^13–15^, with recent evidence demonstrating that epigenetic perturbation alone is sufficient to induce aerobic glycolysis in non-malignant primary cells ^16,17^. At the intersection of glycolysis and OXPHOS, the MPC regulates glucose-derived carbon entry from cytosolic metabolism to mitochondrial metabolism. Alterations in MPC abundance can induce broad metabolic adaptation, including rewiring of biosynthetic pathways. MPC1 and MPC2 expression is markedly low in embryonic stem cells and rises sharply upon differentiation ^18,19^; conversely, reduced MPC expression in proliferative stem and cancer cells sustains aerobic glycolysis, enhances metabolic flexibility, and preserves stem-like features ^20–22^. While complete loss of MPC function is embryonically lethal ^23–25^ partial reduction in mature tissues can profoundly alter cellular physiology without compromising viability.

Canonical Wnt/β-catenin signaling is an established regulator of cellular metabolic state and is strongly activated by truncating *ASXL1* variants in both BOS and myeloid malignancy ^5,11^. β-catenin activation stimulates glucose uptake and transcriptional upregulation of pyruvate carboxylase ^26^. Pyruvate carboxylase catalyzes carboxylation of pyruvate to oxaloacetate, providing carbon to the TCA cycle to enable respiration when other carbon entry points are restricted. Conversely, suppression of Wnt signaling reduces glycolytic dependence through downregulation of PDK1, a central regulator that gates pyruvate entry into the TCA cycle ^27,28^. Wnt activation also induces MYC, which broadly drives glycolysis and mitochondrial remodeling ^29–32^. These observations position Wnt signaling as a key node linking epigenetic perturbation to metabolic reprogramming. Since Wnt signaling both regulates and responds to cellular metabolic state, the causal relationship between Wnt activation and metabolic reprogramming in ASXL1-mutant cells remains unclear.

Germline mutations in BAP1, the catalytic deubiquitinase subunit of the PR-DUB complex and a direct binding partner of ASXL1, have been shown to drive a Warburg-like metabolic state in primary fibroblasts from heterozygous carriers, an effect confirmed by siRNA-mediated BAP1 reduction in vitro ^17,33^. These findings established that PR-DUB complex perturbation is sufficient to induce aerobic glycolysis in non-malignant cells, independent of carcinogenesis. Whether a similar metabolic reprogramming occurs in response to truncating *ASXL1* mutations, which hyperactivate the PR-DUB complex, remains unknown. Moreover, it is unclear whether such metabolic changes contribute to the Wnt dysregulation that we and others have identified in BOS^16,17^.

Here, we investigate the metabolic consequences of truncating ASXL1 mutations in Bohring-Opitz syndrome and define a previously unrecognized connection between epigenetic dysregulation and mitochondrial pyruvate carrier biology. We show that ASXL1 truncation promotes a Warburg-like metabolic state characterized by enhanced glycolytic flux and accumulation of pyruvate and lactate. Mechanistically, truncated ASXL1 and BAP1 exhibit increased occupancy at an H3K4me3-marked regulatory element upstream of MPC2 and selectively reduce MPC1 and MPC2 protein abundance without altering transcript levels. Despite reduced MPC expression, stable isotope tracing demonstrates preserved incorporation of glucose-derived carbon into TCA cycle intermediates, suggesting that *ASXL1*-mutant cells compensate through increased glucose utilization to sustain mitochondrial metabolism.

Pharmacologic MPC inhibition reproduces both the metabolic and Wnt signaling phenotypes of BOS cells, whereas direct Wnt activation alone produces the same metabolic phenotype without affecting MPC protein levels, implicating altered MPC regulation as causally upstream of signaling dysregulation. Together, these findings define a previously unrecognized chromatin-to-metabolism axis through which gain-of-function ASXL1 truncations rewire cellular bioenergetics, and identify MPC as a critical effector linking epigenetic perturbation to metabolic–signaling crosstalk in both a rare developmental syndrome and myeloid malignancy.

## RESULTS

### BOS patient fibroblasts exhibit enhanced glycolytic metabolism and reduced reliance on oxidative phosphorylation

To investigate the metabolic consequences of pathogenic *ASXL1* truncation, we obtained primary dermal fibroblasts from seven individuals with BOS carrying heterozygous truncating or frameshift variants in exons 12–13 of *ASXL1* (**Figure 1A, Table 1**), and eight parental control fibroblast lines. All variants localize to the C-terminal region of ASXL1 and represent a subset of the cohort characterized in our prior multi-omics analysis ^5^. While previous studies established that truncating ASXL1 variants drive broad chromatin remodeling and Wnt signaling dysregulation in patient-derived cells^5^, the metabolic consequences of these mutations had not been investigated. Given prior observations that heterozygous BAP1 germline variants associated a cancer predisposition syndrome perturb the same PR-DUB complex, promote aerobic glycolysis in primary fibroblasts^17,34^, we hypothesized that ASXL1 truncation similarly induces metabolic reprogramming through altered epigenetic regulation.

**Figure 1:**
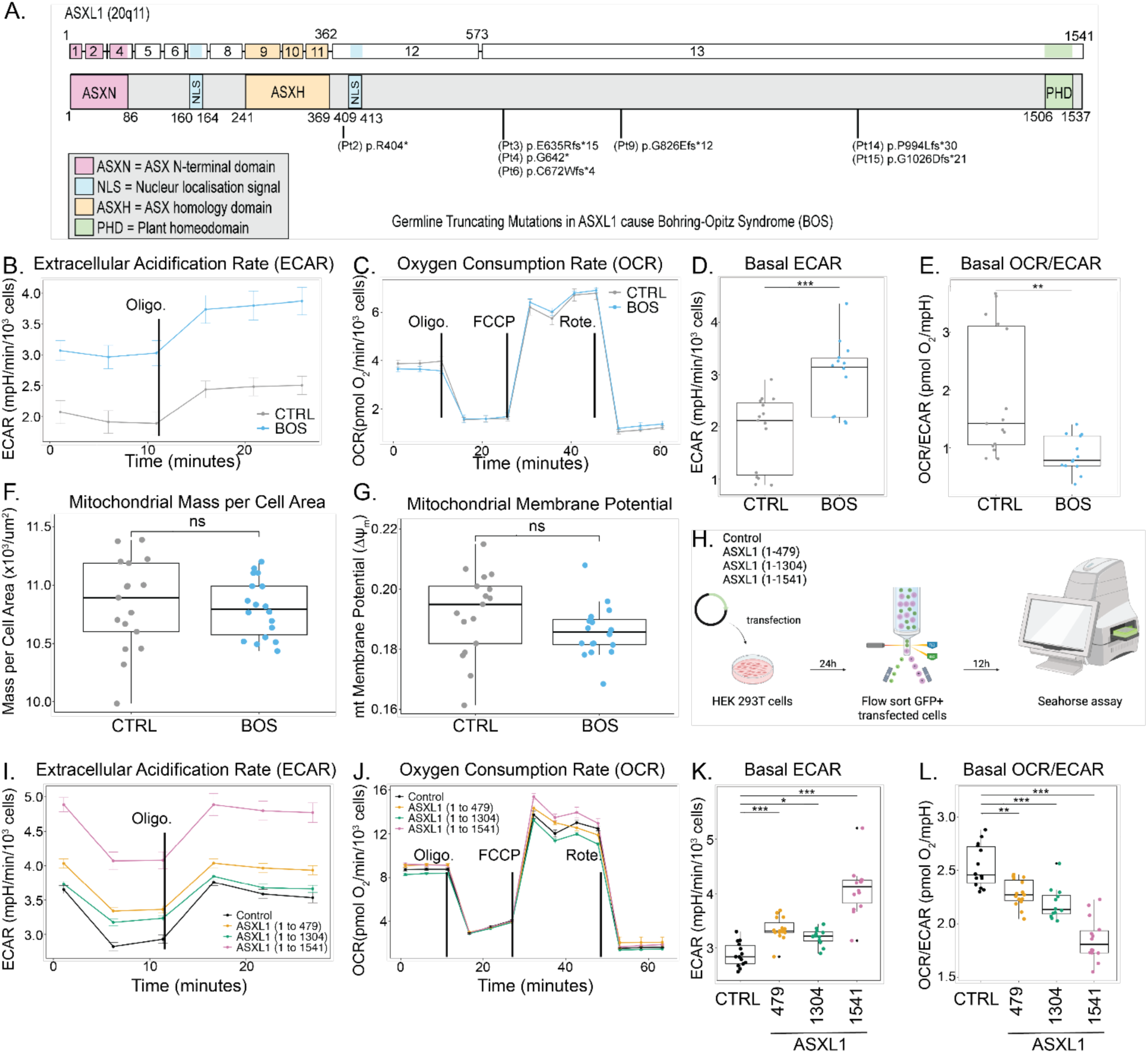
Patient-derived fibroblasts with *ASXL1* truncating variants exhibit enhanced glycolytic metabolism and reduced reliance on oxidative phosphorylation, a metabolic phenotype that is recapitulated by ASXL1 overexpression. **(A)** Schematic of ASXL1 protein domains indicating the location of heterozygous truncating nonsense or frameshift variants identified in seven individuals with Bohring–Opitz syndrome (BOS). All variants localize to exons 12–13 and truncate the C-terminal region of the protein. See Table 1 for full variant details. **(B–E)** Extracellular flux analysis of primary BOS (n = 5) and control (n = 5) fibroblasts. Representative Seahorse traces showing **(B)** extracellular acidification rate (ECAR) and **(C)** oxygen consumption rate (OCR) following sequential injections of oligomycin, FCCP, and rotenone/antimycin A. Quantification of **(D)** basal ECAR and **(E)** basal OCR/ECAR ratio demonstrates significantly elevated levels of glycolysis and reduced mitochondrial reliance in BOS fibroblasts relative to controls. **(F,G)** Mitochondrial morphology and function in BOS and control fibroblasts assessed by fluorescence imaging. No significant differences were observed in **(F)** mitochondrial mass normalized to cell area (MitoTracker Green) or **(G)** mitochondrial membrane potential (TMRE), indicating that altered metabolic flux is independent of gross mitochondrial content or polarization state. **(H)** Schematic of full-length ASXL1 (1–1541) and N-terminal truncation constructs (1–479, 1–1304) used for overexpression studies. GFP-positive cells were isolated by fluorescence-activated cell sorting 24 hours post-transfection and reseeded for downstream analyses. **(I–L)** Extracellular flux analysis of HEK293T cells expressing full-length or truncated ASXL1 constructs relative to empty-vector controls. Representative traces showing **(I)** ECAR and **(J)** OCR. Quantification of **(K)** basal ECAR and **(L)** OCR/ECAR ratio confirms a glycolytic shift across all ASXL1 expression conditions, recapitulating the metabolic phenotype observed in BOS patient-derived fibroblasts. Data are presented as box-and-whisker plots (center line, median; box limits, interquartile range; whiskers, full range) or mean ± s.e.m. ECAR is reported as mpH min⁻¹; OCR as pmol O₂ min⁻¹. Statistical significance was assessed by two-tailed Student’s *t*-test for BOS versus control comparisons or one-way ANOVA with post hoc correction for ASXL1 construct comparisons. Significance levels are indicated as follows: *P < 0.05, **P < 0.01, ***P < 0.001; ns, not significant. See also **Supplemental Figures 1–5, Supplemental Table 1, Extended Data File.**

**Table 1.** Clinical and demographic characteristics of Bohring-Opitz syndrome (BOS) patients and controls. Bohring-Opitz Syndrome (BOS) patients and controls included in our assays are identified with their patient IDs and prior IDs from our previous multi-omics paper ^3^. *ASXL1* mutation, age at collection, and sex of BOS patients are shown. Demographic information for samples where data used were solely from re-analysis of RNA-seq and ATAC-seq in our previous paper are not shown.

| Multi-Omics<br>Paper Pt# | Condition | Sex | Age | ASXL1 Mutation |  | Assays |  |  |  |  |
| --- | --- | --- | --- | --- | --- | --- | --- | --- | --- | --- |
|  |  |  |  | cDNA Change<br>NM_015338.6 | Protein Change<br>Q8IXJ9 | Seahorse | Mitochondrial<br>Analysis | Metabolomics | MPC Protein | mTOR Assays |
| Pt2 | BOS | F | 9 | c.1210C>T | p.R404* |  |  | X |  |  |
| Pt3 | BOS | M | 24 | c.1910_1922del | p.E635Rfs*15 | X | X | X | X | X |
| Pt4 | BOS | M | 8 | c.1924G>T | p.G642* | X |  | X |  | X |
| Pt6 | BOS | F | 15 | c.2013_2014delAG | p.C672Wfs*4 |  |  | X |  |  |
| Pt9 | BOS | F | 3 | c.2477delG | p.G826Efs*12 | X | X |  | X | X |
| Pt14 | BOS | F | 5 | c.2981delC | p.P994Lfs*30 | X | X |  | X | X |
| Pt15 | BOS | F | 7 | c.3077delG | p.G1026Dfs*21 | X |  | X |  |  |
| Ctrl1 | Control | F | 37 |  |  |  |  |  | X |  |
| Ctrl2 | Control | M | 38 |  |  | X |  | X |  |  |
| Ctrl3 | Control | F | 40 |  |  | X |  | X |  | X |
| Ctrl5 | Control | F | 41 |  |  | X | X |  | X | X |
| Ctrl50 | Control | F | 56 |  |  | X | X |  |  |  |
| Ctrl51 | Control | M | 55 |  |  | X | X | X | X | X |
| Ctrl52 | Control | F | 45 |  |  |  |  | X | X | X |
| Ctrl53 | Control | M | 44 |  |  |  |  | X | X | X |

To determine whether *ASXL1* truncating variants alter cellular metabolism, we first assessed cellular bioenergetics in patient-derived fibroblasts. BOS fibroblasts exhibited a significant increase in extracellular acidification rate (ECAR, mean 3.00 mpH/min/10^3^ cells vs 1.90 mpH/min/10^3^ cells control, p = 1.01E-04), indicating elevated glycolytic flux, accompanied by a modest but significant reduction in basal oxygen consumption rate (OCR, mean 2.45 pmol/min BOS vs 2.93 pmol/min control, p = 4.23E-02), resulting in a markedly reduced OCR/ECAR ratio (**Figures 1B - D, Supplemental Table 1)**. This metabolic shift demonstrates preferential reliance on glycolysis over oxidative metabolism and was reproducible across independent experimental replicates (**Figure 1E**).

To further characterize the bioenergetic shift, we quantified the relative contributions of glycolysis and oxidative phosphorylation to total cellular ATP production. BOS fibroblasts generated substantially reduced total ATP via oxidative phosphorylation (mean 35.37% ± 10.95%) compared to controls (mean 52.77% ± 16.93%) (p=1.16E-03), indicating increased reliance on glycolytic ATP generation (**Supplemental Figure 1**). Notably, maximal respiratory capacity (maximal OCR) remained largely intact in BOS cells, indicating that mitochondrial oxidative metabolism is not intrinsically impaired. Rather, BOS cells retain the capacity to engage oxidative phosphorylation when required, but do so at a chronically reduced set point under basal conditions.

Consistent with a functional rather than structural mitochondrial defect, mitochondrial mass and membrane potential were unchanged between BOS and control fibroblasts (**Figures 1F – G, Supplemental Figure 2)**, indicating that the metabolic reprogramming occurs independently of gross mitochondrial content or polarization state. Together, these data demonstrate that truncating *ASXL1* variants drive a reproducible and selective reprogramming of cellular energy metabolism toward aerobic glycolysis, characterized by increased glycolytic flux despite preserved mitochondrial infrastructure. This pattern is most consistent with a regulated metabolic shift rather than overt mitochondrial dysfunction.

### ASXL1 overexpression and truncation recapitulate metabolic rewiring in HEK293T cells

To determine whether ASXL1 perturbation is sufficient to drive this metabolic phenotype independently of the patient-specific genetic background of BOS fibroblasts, we overexpressed GFP-tagged full-length ASXL1 (1–1541) and N-terminal truncation constructs (1–479 and 1–1304) in HEK293T cells^35^ (**Figure 1H**). GFP-positive cells were isolated by fluorescence-activated cell sorting 24 hours post-transfection to ensure analysis of a pure expressing population **(Supplemental Figures 3 - 4)**, and extracellular flux analysis was performed 24 hours after reseeding.

Expression of either truncating ASXL1 constructs significantly increased ECAR (p < 0.001) relative to empty-vector controls, recapitulating the metabolic profile observed in BOS patient-derived fibroblasts (Figures 1I – L). In contrast, OCR was largely unchanged, suggesting that increased glycolytic activity can occur in the absence of substantial reductions in mitochondrial respiration. Consistent with these findings, cells expressing truncating *ASXL1* variants exhibited increased reliance on glycolytic ATP production **(Supplemental Figure 5)**.

Notably, overexpression of full-length ASXL1 produced a similar metabolic shift, indicating that the phenotype is sensitive to ASXL1 dosage and activity rather than being uniquely dependent on truncation-derived protein products. The consistency of this metabolic shift across two truncation constructs representing distinct C-terminal endpoints (1–479 and 1–1304) and full-length ASXL1 suggests that the metabolic effect is driven by the N-terminal region common to all constructs rather than by truncation-specific C-terminal sequences. Together with the patient fibroblast data, these findings demonstrate that ASXL1 perturbation is sufficient to drive a Warburg-like metabolic state across distinct cellular contexts, supporting a gain-of-function mechanism for truncating ASXL1 variants in metabolic reprogramming.

#### Metabolomics and isotope tracing show increased reliance on glycolysis in BOS fibroblasts

To define the broader metabolic consequences of *ASXL1* truncation beyond extracellular flux measurements, we performed metabolomic profiling of BOS (n = 4) and control (n = 6) fibroblasts **(Supplemental Table 2)**. Given the increased glycolytic utilization in BOS fibroblasts, which is a prominent finding in pluripotent cells and in tumors^36–39^, we asked whether this was a singular derangement or if other metabolic pathways were affected. We performed targeted and untargeted U-¹³C-glucose and U-¹³C-glutamine tracing to understand how carbons moved through the cell.

In our untargeted metabolomics study on control and BOS fibroblasts, we use orthogonal partial least squares discriminant analysis (OPLS-DA) to cluster our samples and these demonstrated a clear separation along the predictive T-score axis, with BOS and control fibroblasts clustering into non-overlapping groups, indicating a distinct metabolic profile associated with the *ASXL1* truncating variant (**Figure 2A**). Pathway-level enrichment analysis using Mummichog 2.0 identified a significant enrichment of glycolysis/gluconeogenesis (enrichment factor 2.05, -log10 p-value=2.16) and pyruvate metabolism (enrichment factor 2.34, -log10 p-value=1.96) (**Figure 2B, Supplemental Table 3**), consistent with increased glycolysis without impairment to pyruvate utilization. Gene ontology analysis independently identified the Warburg effect as the most strongly enriched metabolic phenotype (fold enrichment = 6.76, FDR = 0.0185), with additional enrichment of glycolysis (fold enrichment = 10.07, FDR = 0.0638) and gluconeogenesis (fold enrichment = 9.35, FDR = 0.0185) (**Figure 2C, Supplemental Table 4)**.

**Figure 2:**
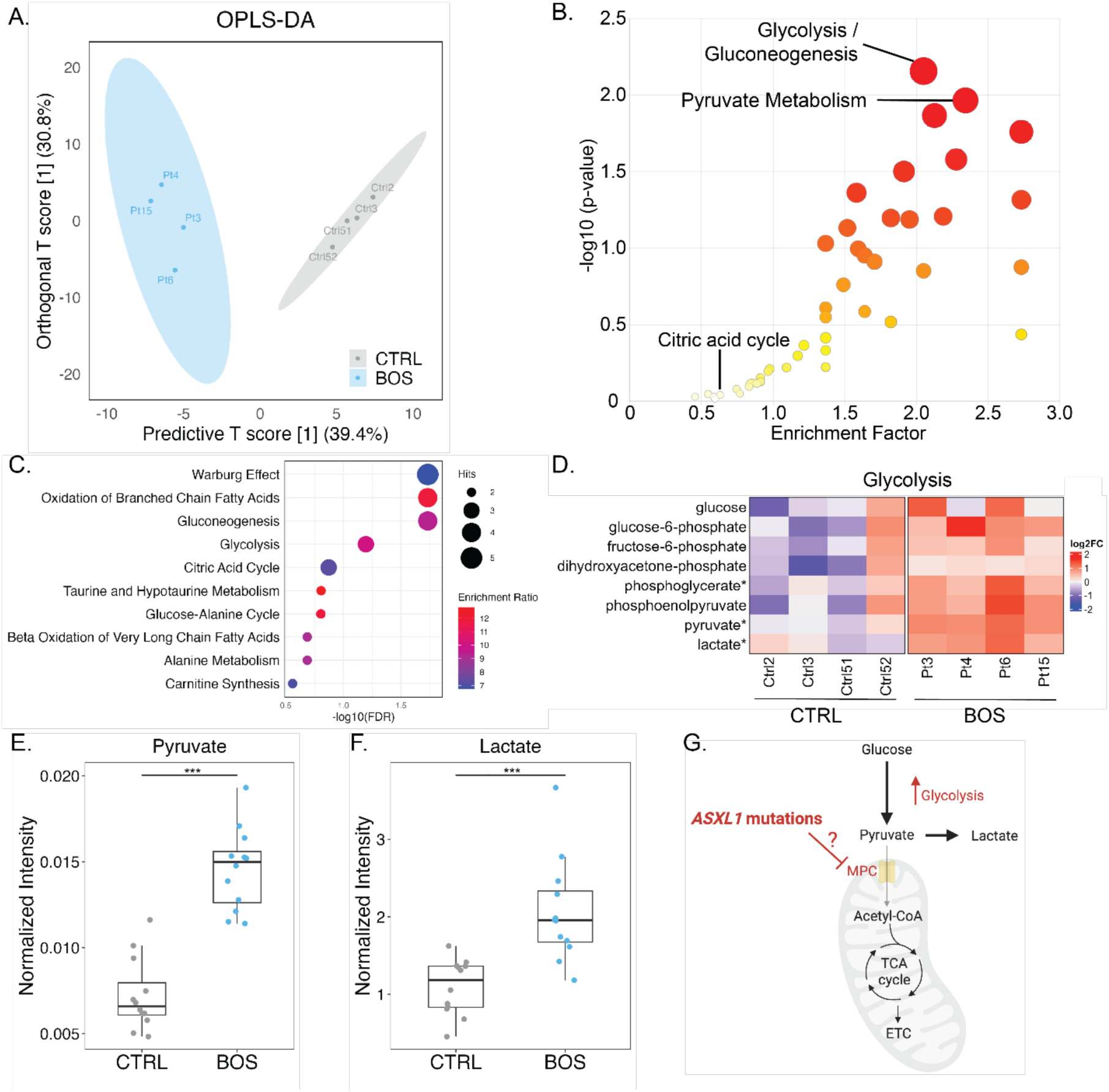
Untargeted metabolomics reveal increased glycolytic throughput and pyruvate accumulation in BOS patient-derived fibroblasts. **(A)** OPLS-DA score plot of untargeted metabolomics from BOS (n = 4) and control fibroblasts (n = 4), demonstrating clear separation of metabolic profiles. **(B)** Pathway enrichment analysis (Mummichog 2.0) identifies glycolysis/gluconeogenesis, pyruvate metabolism, and TCA cycle pathways as significantly enriched. **(C)** Gene ontology analysis highlights the Warburg effect as the most significantly enriched metabolic phenotype. **(D)** Heatmap of glycolytic pathway showing increased abundance in BOS fibroblasts (n = 4) relative to controls (p < 0.05). **(E,F)** Relative abundance of pyruvate **(E)** and lactate **(F)** is significantly increased in BOS fibroblasts. **(G)** Proposed model of metabolic rewiring associated with ASXL1 truncating mutations. *Data are shown as box-and-whisker plots (median, interquartile range, and range) or mean ± s.e.m. Statistical significance was determined using two-tailed Student’s t-tests with false discovery rate correction where applicable*. *Significance levels are indicated as follows: *P < 0.05, **P < 0.01, ***P < 0.001; ns, not significant*. **See also** Supplemental Figures 6-11, Supplemental Tables 2-4, Extended Data File.

We next performed targeted U-¹³C-glucose tracing for control and BOS fibroblasts. After 6 hours in tracing media, glycolytic intermediates were broadly elevated in BOS fibroblasts (**Figure 2D, Supplemental Figure 6B)**, accompanied by significantly increased intracellular pyruvate (p = 4.03 × 10⁻⁴) and lactate (p = 6.32 × 10⁻⁸) (**Figures 2E – F, Supplemental Figures 6D - E)**, consistent with the extracellular acidification data and indicating intracellular accumulation of glycolytic end products.

To distinguish between increased glycolytic throughput and impairment of specific enzymatic steps, we performed stable isotope tracing using uniformly labeled U-¹³C-glucose and U-¹³C-glutamine. Isotopologue distribution analysis at 24 hours revealed remarkably similar labeling patterns between BOS and control fibroblasts, with approximately 70% M+6 and 30% M+0 in both groups, and pyruvate showed complete M+3 labeling from U-¹³C-glucose in both conditions **(Supplemental Figure 7A - B)**. Likewise, the relative contribution of glucose- and glutamine-derived carbon to lactate pools were similar between control and BOS fibroblasts **(Supplemental Figure 7C)** indicating that the enzymatic steps of glycolysis are intact, and that the elevated glycolytic metabolite pools observed in BOS fibroblasts likely arise from increased pathway flux.

Extended isotope tracing at 6, 24, and 72 hours after media change supported these findings. BOS fibroblasts consistently exhibited increased pool sizes of glycolytic intermediates and end products across all time points, while isotopologue distributions remained comparable to controls **(Supplemental Figures 8 - 10)**. Strikingly, the difference in glycolytic metabolite abundance was most pronounced at the 6-hour time point since media change and diminished progressively suggesting a dependence on fresh media or accumulation of an inhibitory molecule that diminish overall effect.

TCA cycle metabolites were maintained or modestly elevated in BOS fibroblasts relative to controls **(Supplemental Figures 8 - 10)**. To investigate potential compensatory mechanisms, we examined metabolic pathways contributing carbon to mitochondrial metabolism. Metabolite set enrichment analysis identified significant alterations in glutamate metabolism, the malate–aspartate shuttle, carnitine synthesis, and the pentose phosphate pathway **(Supplemental Figure 11A)**. Consistent with these findings, BOS fibroblasts exhibited increased abundance of aspartate and glutamate together with increased incorporation of glucose-derived carbon into both metabolites **(Supplemental Figure 11B - C)**, suggesting that BOS fibroblasts predominantly maintain TCA cycle intermediates through pyruvate utilization.

Additional metabolic alterations were observed across pathways involved in redox homeostasis, lipid metabolism, and glycolysis branch pathways. BOS fibroblasts exhibited significantly increased NAD⁺ abundance, consistent with altered cellular redox metabolism **(Supplemental Figure 11D)**. Acetyl-carnitine levels were also elevated in BOS fibroblasts despite similar isotopologue distributions between BOS and control fibroblasts, suggesting additional utilization of alternative mitochondrial carbon sources **(Supplemental Figure 11E)**. Furthermore, metabolites of the polyol pathway, including sorbitol, were significantly increased while retaining predominantly glucose-derived (M+6) labeling **(Supplemental Figure 11F)**. Enrichment of these pathways suggests that excess glucose may be diverted into overflow metabolic pathways, and that truncating *ASXL1* mutations may drive broader compensatory metabolic rewiring.

Together, these data establish that *ASXL1* truncation promotes increased glucose utilization, accumulation of glycolytic intermediates (**Figure 1**), and accumulation of pyruvate and lactate in BOS fibroblasts (**Figure 2E - F**). Despite this pronounced glycolytic phenotype, isotope tracing revealed preserved incorporation of glucose-derived carbon into downstream mitochondrial metabolites, indicating that mitochondrial carbon supply remains largely maintained. These findings suggest that ASXL1-mutant cells adapt to metabolic perturbation by increasing glycolytic throughput while sustaining TCA cycle metabolism. Interestingly, the BOS clinical phenotype includes abundant hair growth, which has been associated with mitochondrial pyruvate carrier (MPC) inhibition^40^, so we next asked whether the MPC is directly dysregulated in the presence of truncating *ASXL1* mutations (**Figure 2G**).

### *ASXL1* truncating variants expression reduce mitochondrial pyruvate carrier protein levels across cellular models

MPC functions as an obligate heterodimer of MPC1 and MPC2, and loss of either subunit destabilizes the complex and alters mitochondrial pyruvate import ^41^. Reduced MPC abundance is associated with increased reliance on glycolysis for ATP production, even in the presence of oxygen, similar to what is observed in Warburg effect ^42^. Because BOS fibroblasts exhibited enhanced glycolytic flux together with accumulation of pyruvate and lactate, we asked whether truncating ASXL1 variants alter expression of the MPC complex. We therefore examined MPC protein levels in patient-derived fibroblasts and complementary cellular models.

Western blot analysis of BOS patient-derived fibroblasts (n=3 biological replicates) compared to controls (n=4 biological replicates) demonstrated a significant reduction in both MPC1 and MPC2 protein levels (**Figure 3A**). Quantification normalized to β-actin, revealed a fold change of 0.409 for MPC1 (p = 4.31E-6, Welch two sample t-test) and 0.659 for MPC2 (p = 2.31E-5) relative to controls (**Figure 3B, Supplemental Table 5**), indicating that both subunits of the MPC heterodimer are reduced in BOS patient-derived cells.

**Figure 3:**
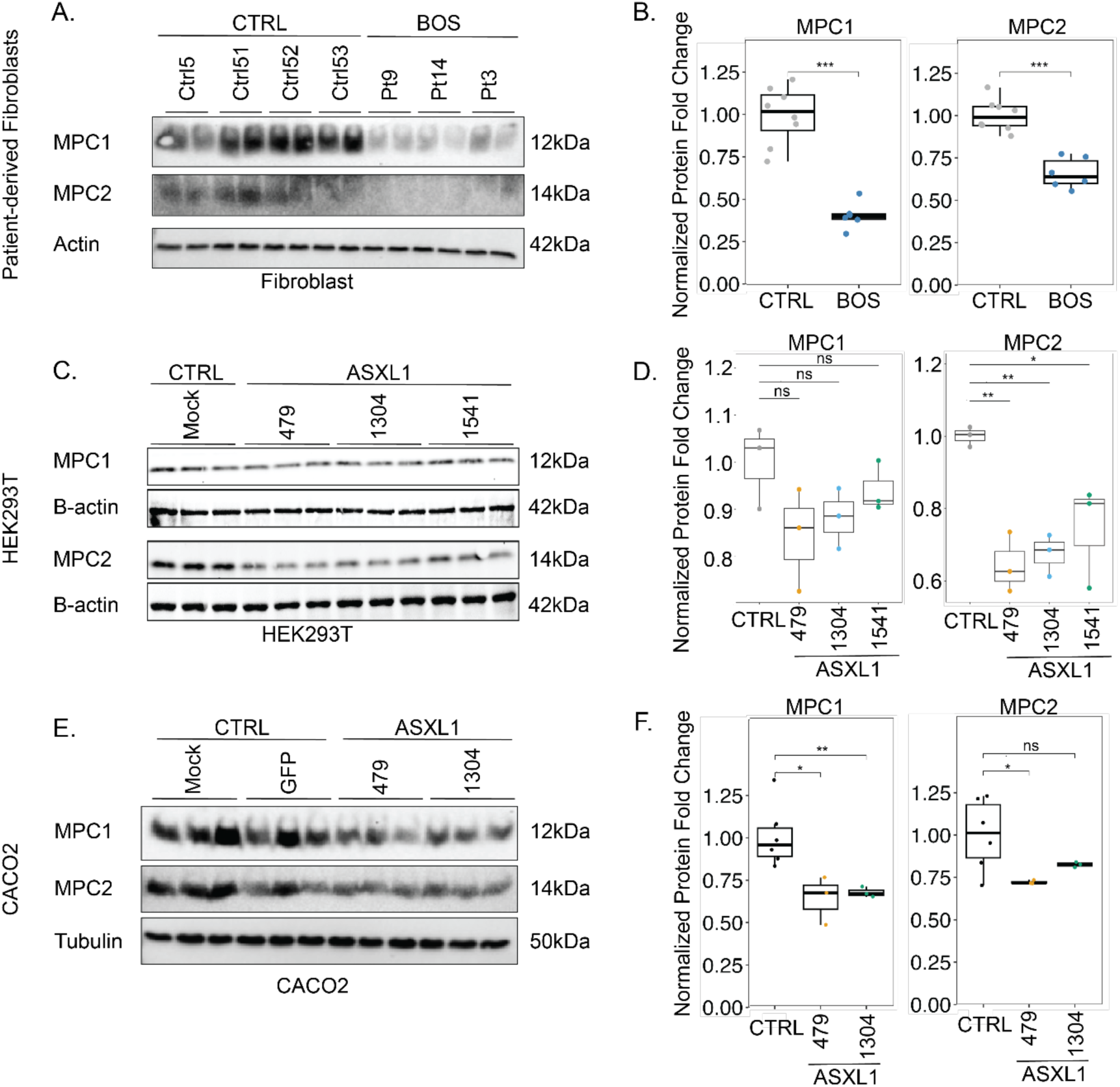
A*S*XL1 truncating variants reduce mitochondrial pyruvate carrier protein levels across cellular models. **(A)** Representative immunoblots of MPC1 and MPC2 in primary control and BOS fibroblasts; â-actin serves as a loading control. **(B)** Quantification of MPC1 and MPC2 protein levels from (**A**) normalized to β-actin. Both proteins are significantly reduced in BOS fibroblasts. **(C)** Representative immunoblots of MPC1 and MPC2 in flow sorted HEK293T cells overexpressing GFP-tagged full-length or truncated ASXL1 constructs. **(D)** Quantification of MPC protein levels from (**C)**, showing significant reduction of MPC2 across all ASXL1 constructs. **(E)** Representative immunoblots **(F)** and quantification of MPC1 and MPC2 in Caco-2 cells expressing ASXL1 truncation constructs. *Data are shown as box-and-whisker plots (median, interquartile range, and range) or mean ± s.e.m. Statistical significance was assessed using two-tailed Student’s t-tests. Significance levels are indicated as follows: *P < 0.05, **P < 0.01, ***P < 0.001; ns, not significant*.

To determine whether reduced MPC abundance is directly attributable to ASXL1 perturbation rather than patient-specific genetic background, we examined MPC protein levels in HEK293T cells expressing full-length or truncated ASXL1 constructs. Expression of either truncating constructs (1–479 and 1–1304) significantly reduced MPC1 and MPC2 protein levels relative to empty-vector controls, and full-length ASXL1 overexpression produced a similar reduction (**Figures 3C–D, Supplemental Figure 12, Supplemental Table 5)**, consistent with the dosage-sensitive metabolic phenotype observed in extracellular flux analyses. Interestingly, in fluorescence-sorted GFP-positive cells analyzed using whole-cell lysates, MPC2 showed the most consistent reduction, whereas in unsorted cells using cytoplasmic extracts, MPC1 exhibited the larger effect.

This reduction in MPC levels was independently confirmed in Caco-2 cell lines, an immortalized colorectal cancer cell line, expressing truncating ASXL1 constructs (**Figures 3E–F, Supplemental Figure 13, Supplemental Table 5)**. Together, these data establish that *ASXL1* perturbation drives a selective and reproducible reduction of MPC protein levels across multiple differentiated cellular contexts, defining reduced MPC abundance as a conserved molecular consequence of ASXL1 dysregulation. Notably, this reduction occurs in the setting of enhanced glycolytic flux but preserved incorporation of glucose-derived carbon into TCA cycle intermediates. The mechanism underlying this protein-level reduction is examined in **Figure 5**.

### Pharmacologic MPC inhibition recapitulates the BOS metabolic and Wnt signaling phenotype and establishes reduced MPC abundance as upstream of Wnt activation

We next sought to define the causal hierarchy between reduced MPC abundance, metabolic reprogramming, and dysregulation of Wnt-signaling in *ASXL1*-mutant cells. We first ask whether reduced MPC abundance was sufficient to drive the metabolic changes observed in BOS. Given our previous work identifying dysregulated Wnt signaling in ASXL1, we also asked whether Wnt-activation functions upstream or downstream of these metabolic alterations.

To validate that the metabolic changes are driven primarily by MPC loss, we pharmacologically inhibited the mitochondrial pyruvate carrier using UK5099 ^43,44^ and assessed metabolic flux and downstream signaling. UK5099 treatment produced a dose-dependent increase in AXIN2 protein levels accompanied by altered β-catenin abundance, indicating activation of canonical Wnt signaling (**Figure 4A, Supplemental Table 6)**.

**Figure 4:**
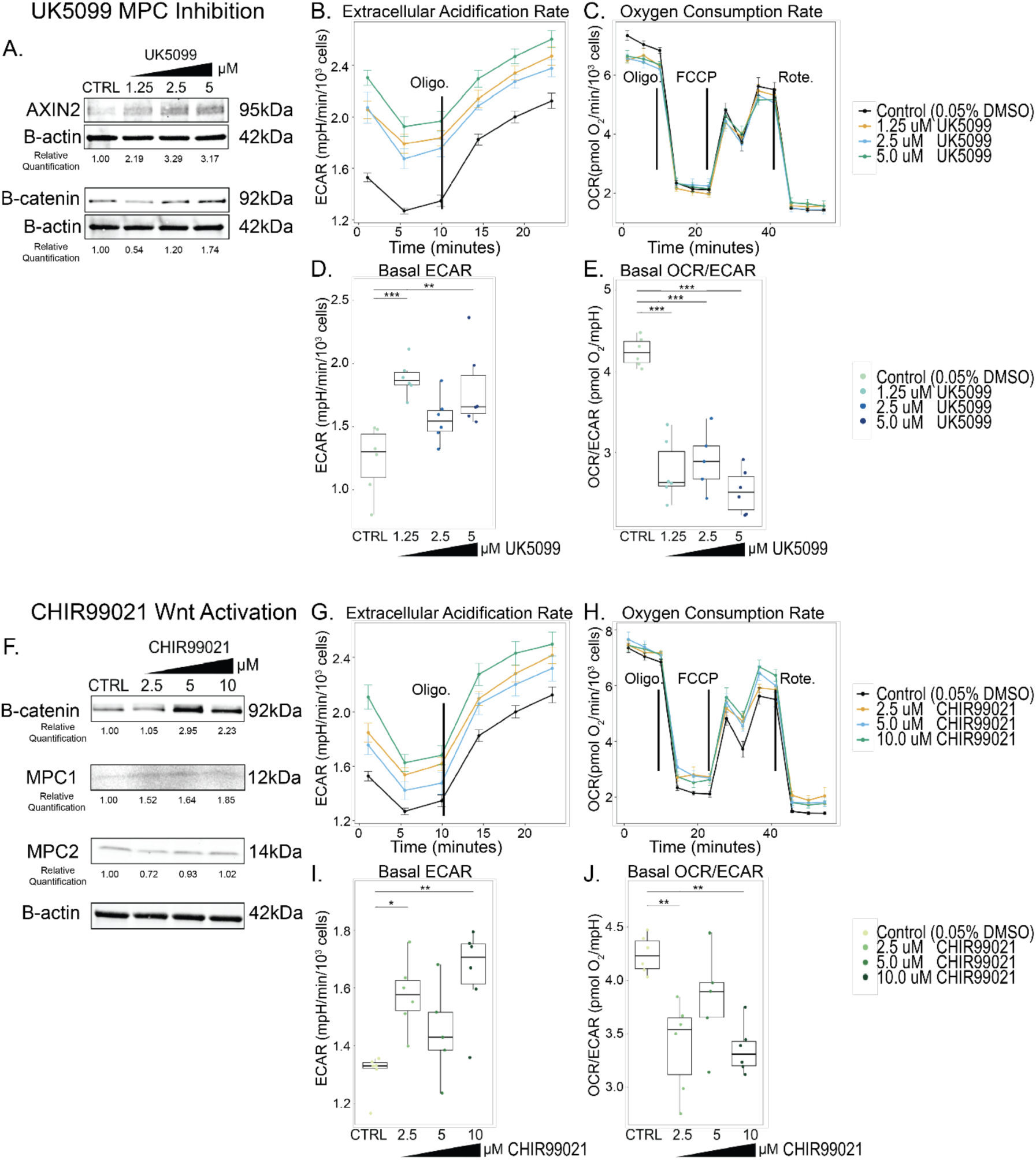
Pharmacologic MPC inhibition recapitulates the BOS metabolic and Wnt signaling phenotype, and establishes reduced MPC abundance as upstream of Wnt activation. (A) Representative immunoblots of AXIN2 and â-catenin in cells treated with increasing concentrations of the MPC inhibitor UK5099 (0, 1.25, 2.5, and 5 μM) for 24 hours. AXIN2 expression increases in a dose-dependent manner following MPC inhibition. **(B - E)** Seahorse extracellular flux analysis following MPC inhibition with UK5099 demonstrates (B) increased extracellular acidification rate (ECAR) and (**C**) reduced oxygen consumption rate (OCR) (p = 0.0232). **(D)** Basal ECAR is significantly increased, and **(E)** basal OCR/ECAR ratio is significantly decreased with increasing concentrations of UK5099. **(F)** Representative immunoblots of â-catenin, MPC1 and MPC2 in cells treated with increasing concentrations of the Wnt agonist CHIR99021 (0, 2.5, 5, and 10 μM), confirm dose-dependent activation of Wnt signaling, and MPC1 levels, while MPC2 protein abundance remains unchanged. **(G - H)** Seahorse extracellular flux analysis following Wnt activation with CHIR99021 shows (**G**) Basal ECAR is significantly increased with (**H**) no significant change in OCR. **(I)** Basal ECAR is significantly increased, and **(J)** basal OCR/ECAR ratio is significantly decreased with increasing concentrations of CHIR99021. Data are shown as box-and-whisker plots (median, interquartile range, and range) or mean ± s.e.m. ECAR is reported as mpH min⁻¹ and OCR as pmol O₂ min⁻¹. Statistical significance was assessed using two-tailed Student’s t-tests or one-way ANOVA with post hoc correction. Significance levels are indicated as follows: *P < 0.05, **P < 0.01, ***P < 0.001; ns, not significant.

Extracellular flux analysis demonstrated that UK5099 induced a dose-dependent reduction in OCR together with increased ECAR (**Figures 4B - E, Supplemental Table 1)**, recapitulating the metabolic profile observed in BOS patient-derived fibroblasts and ASXL1-expressing cell models. These data establish that reduced MPC abundance is sufficient to drive both the Warburg-like metabolic state and the Wnt signaling activation that characterize BOS cells.

To address the question as to whether Wnt activation is sufficient to drive the metabolic changes, we treated cells with the GSK3β inhibitor CHIR99021, which activates canonical Wnt signaling through stabilization of β-catenin. CHIR99021 induced robust dose-dependent accumulation of β-catenin, confirming effective Wnt pathway activation (**Figure 4F, Supplemental Table 6)**. Prior studies had observed that MPC expression was negatively correlated with expression of Wnt/B-catenin target genes ^22^, however, the mechanism and direction of causality have not been established. We found that MPC1 levels increased while MPC2 protein abundance remained unchanged across increasing CHIR99021 concentrations (**Figure 4F**), demonstrating that Wnt activation does not reduce MPC levels in our studies.

Extracellular flux analysis revealed that CHIR99021 increased glycolytic flux, as reflected by significantly elevated ECAR and a reduced OCR/ECAR ratio (**Figures 4G–J**), consistent with the established role of canonical Wnt/β-catenin signaling in promoting aerobic glycolysis. However, OCR was found to have a modest, though not statistically significant, increase as a result of Wnt activation (**Figure 4G**), in contrast to the mild OCR reduction produced by UK5099 treatment (p-value = 0.232) (**Figure 4C**). This distinction demonstrates that Wnt activation can independently promote glycolysis but does not act through MPC complex downregulation.

These findings were reproduced in independent biological replicate experiments **(Supplemental Figure 14)**. These data support a model in which ASXL1-truncation-dependent loss of MPC protein reduces mitochondrial pyruvate import, driving glycolytic reprogramming, and secondary activation of Wnt signaling. While Wnt activation promotes glycolytic metabolism, it does so without altering MPC abundance. Thus, mitochondrial pyruvate transport acts upstream of Wnt activation, and reduced MPC abundance may be the primary driver of the adaptive metabolic reprogramming seen in BOS cells.

### ASXL1 truncation alters nutrient-sensing responses independently of canonical ATF4–ASNS transcriptional regulation

The metabolic remodeling in BOS fibroblasts prompted examination of nutrient sensing pathways. BOS fibroblasts exhibited reduced pS6K under amino acid-replete conditions and progressive mTOR pathway impairment following non-essential amino acid deprivation at 16, 48, and 72 hours, with reduced ATF4 protein during early amino acid stress **(Supplemental Figure 15A)**. L-asparagine partially restored pS6K signaling, consistent with asparagine coupling mitochondrial respiration to mTOR–ATF4 signaling ^45^. Preserved H3K4me3 at the *ATF4* promoter and unchanged *ATF4* and *ASNS* transcript levels confirmed that impaired nutrient sensing arises downstream of mitochondrial metabolic signaling rather than epigenetic silencing **(Supplemental Figure 16)**.

### Truncated ASXL1 and BAP1 co-occupy an H3K4me3-marked regulatory element upstream of *MPC2*

We interrogated publicly available ChIP-seq datasets generated in HEK293T cells expressing GFP-tagged full-length or truncated ASXL1 constructs (GSE302969), complemented by matched histone modification profiling (ENCFF439DDQ for H3K4me3; ENCFF885SUR for H3K27Ac)^46,47^. These datasets, generated independently from our patient-derived and transfection models, allowed direct comparison of full-length and truncated ASXL1 chromatin occupancy and provided an independent framework for investigating potential regulatory interactions at the MPC locus.

At the *MPC2* locus, both full-length and truncated ASXL1 show occupancy at a regulatory element within *MPC2* intron 1, between the transcription start site in exon 1 and the translation start codon in exon 2 (**Figure 5A**). Full-length ASXL1 exhibits a discrete, focused peak at this element, while truncated ASXL1 exhibits a broader, less focused occupancy profile extending beyond full-length boundaries — consistent with loss of C-terminal targeting specificity. BAP1 shows co-occupancy confirming PR-DUB complex engagement. BRD4 occupies the same element with a focused peak unchanged between full-length and truncated ASXL1 conditions. This dissociation — ASXL1 occupancy broadens while BRD4 remains focused — indicates loss of positional targeting precision rather than gain of transcriptional activity, consistent with aberrant PR-DUB spreading. The locus is marked by H3K4me3, consistent with a promoter-associated regulatory element or putative alternative TSS. The *MPC1* locus showed no discrete intronic peak for ASXL1, BAP1, or BRD4 (**Figure 5B**), with secondary MPC1 loss reflecting heterodimer destabilization ^48^. ATAC-seq profiling of BOS fibroblasts confirmed two accessible chromatin regions: a preserved double-peak signature at the canonical *MPC2* promoter ruling out promoter silencing, and a second region within intron 1 partially overlapping the ChIP-seq peaks with subtle increased accessibility in BOS fibroblasts, confirming this intronic element is an active regulatory site in patient-derived cells **(Supplemental Figure 25)**. No β-catenin occupancy differences were observed at *MPC1* or *MPC2* following Wnt stimulation **(Supplemental Figure 17)**, ruling out a direct transcriptional Wnt-to-MPC connection.

**Figure 5:**
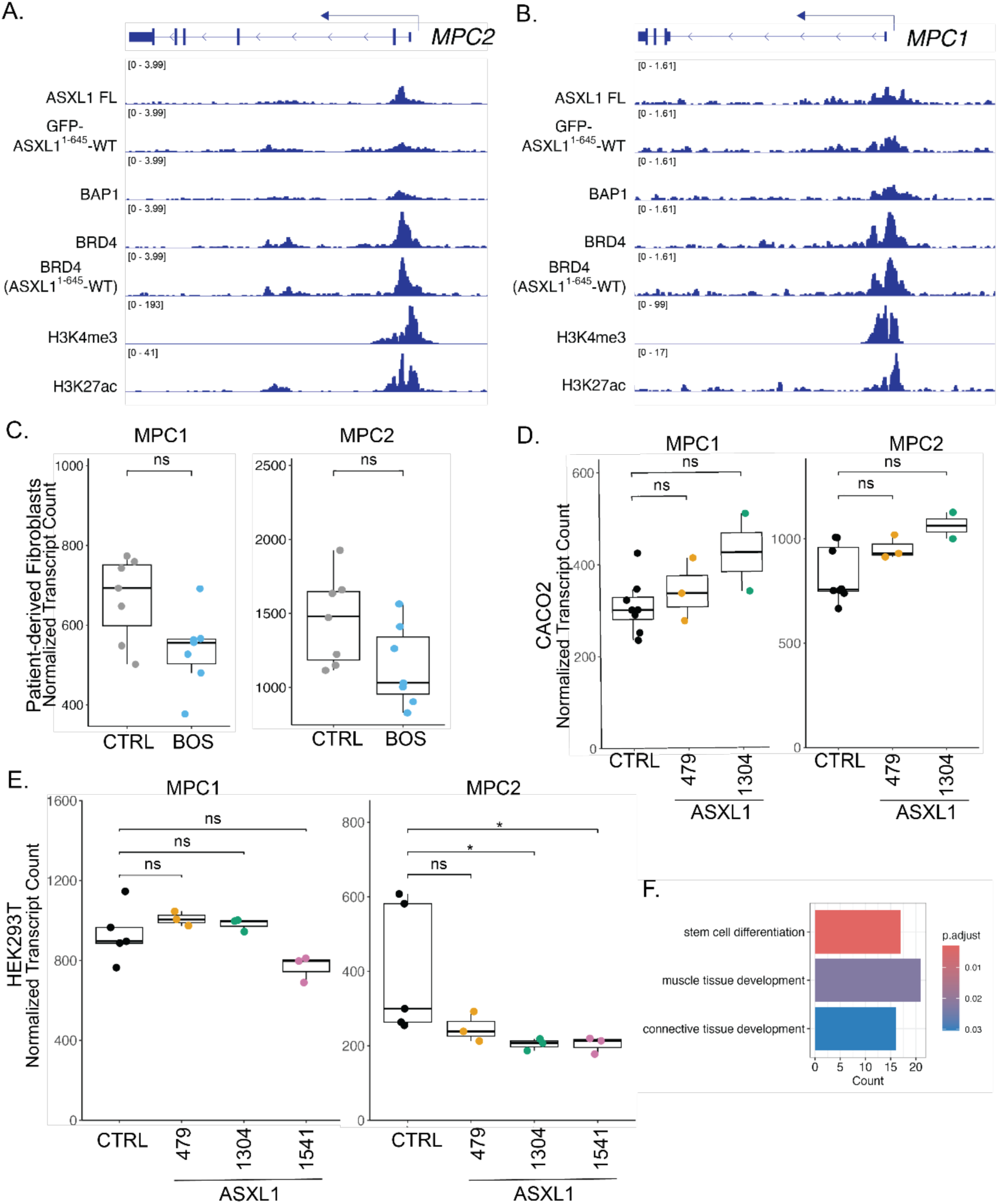
ASXL1 regulates MPC2 expression through binding of the MPC2 promoter. **(A)** Genome browser tracks at the *MPC2* locus showing ChIP-seq occupancy of full-length ASXL1-FL, a truncated form GFP-ASXL1^1–645^, BAP1, BRD4, and BRD4 cotransfected with GFP-ASXL1^1–645(GSE302969)^, and alongside histone modification marks H3K4me3 (ENCFF439DDQ) and H3K27Ac (ENCFF885SUR) in HEK293T cells. A regulatory element within MPC2 intron 1, between the transcription start site in exon 1 and the translation start site in exon 2, shows discrete focused occupancy of full-length ASXL1 and broader, less specific occupancy of truncated GFP-ASXL1^1–645^ at the same locus, The altered occupancy profile of truncated ASXL1 is consistent with loss of C-terminal targeting specificity and aberrant spreading of PR-DUB complex activity across the intronic regulatory region. **(B)** Genome browser tracks at the *MPC1* locus using the same ChIP-seq datasets, shown as a specificity control. No discrete upstream regulatory peak is observed at the *MPC1* locus, consistent with the primary reduction of MPC2 at the chromatin level and secondary MPC1 loss through impaired MPC1/MPC2 heterodimer stability. **(C)** Normalized RNA sequencing transcript counts for *MPC1* and *MPC2* in BOS patient-derived fibroblasts and control fibroblasts. Neither transcript demonstrates significant differential expression between groups. **(D)** Normalized RNA sequencing transcript counts for *MPC1* and *MPC2* in Caco-2 cells expressing ASXL1 truncation constructs (1–479 and 1–1304) relative to controls. No significant changes in *MPC1* or *MPC2* transcript abundance were observed. **(E)** Normalized RNA sequencing transcript counts for *MPC1* and *MPC2* in HEK293T cells expressing ASXL1 constructs (1–479, 1–1304, and 1–1541) relative to controls. Modest reductions in *MPC2* transcript abundance were observed in cells expressing ASXL1 1–1304 (log₂FC = −0.322, adjusted P = 0.048) and ASXL1 1–1541 (log₂FC = −0.346, adjusted P = 0.024), whereas no significant change was observed with ASXL1 1–479. *MPC1* transcript abundance remained unchanged across conditions. **(F)** Gene ontology analysis of differentially expressed genes identified in HEK293T cells expressing ASXL1 constructs relative to controls. Significant pathways include stem cell differentiation, connective tissue development, and muscle tissue development. *Data are shown as box-and-whisker plots (median, interquartile range, and range) or mean ± s.e.m. Statistical significance was assessed using two-tailed Student’s t-tests. Significance levels are indicated as follows: *P < 0.05, **P < 0.01, ***P < 0.001; ns, not significant*.

### Modest *MPC2* transcriptional changes are amplified at the protein level through post-transcriptional mechanisms

MPC1 and MPC2 transcript levels were unchanged in BOS fibroblasts and Caco-2 cells expressing truncating ASXL1 constructs (**Figures 5C–D**). In HEK293T cells, modest but significant *MPC2* reductions were observed with ASXL1 1–1304 (log₂FC = −0.322, adjusted *P* = 0.048) and 1–1541 (log₂FC = −0.346, adjusted *P* = 0.024) but not 1–479 (**Figure 5E**). These modest changes are substantially smaller than protein-level reductions across all systems, indicating post-transcriptional amplification. Broader transcriptomic analysis identified 350 DEGs in HEK293T cells and 704 in Caco-2 cells, with GO enrichment identifying proteasome-mediated ubiquitin-dependent catabolic processes in full-length ASXL1-expressing HEK293T cells and respiratory chain complex assembly pathways in Caco-2 cells (**Figure 5F, Supplemental Figures 18–22)**. Cross-model analysis identified 439 shared DEGs enriched in RNA processing pathways, and no PDK family member showed significant transcriptional change **(Supplemental Figure 23)**. Together, these findings support a chromatin-to-protein axis in which aberrant ASXL1/BAP1 co-occupancy at the H3K4me3-marked intronic *MPC2* element — characterized by broadened ASXL1 occupancy dissociated from stable BRD4 positioning — initiates modest transcriptional dysregulation amplified at the protein level through post-transcriptional mechanisms.

## DISCUSSION

Our findings establish that truncating *ASXL1* mutations, which hyperactivate the PR-DUB complex through aberrant BAP1 stabilization, produce a coherent and reproducible metabolic phenotype traceable to a single functional bottleneck: impaired mitochondrial pyruvate import. Across patient-derived fibroblasts, heterologous overexpression models, and pharmacologic perturbation, evidence converges on the MPC complex as a central mechanistic node directly regulated by ASXL1 truncation. Pharmacologic MPC inhibition recapitulates not only the metabolic profile of BOS cells but also the Wnt signaling activation previously identified in patient-derived tissues^5^, establishing MPC loss as causally upstream rather than merely correlative. The negative result is equally informative: direct Wnt activation contributes to increased glycolytic flux without suppressing mitochondrial respiration or reducing MPC levels, ruling out Wnt dysregulation as the primary driver of the metabolic phenotype. Our work establishes a causal hierarchy of chromatin dysregulation: MPC loss leads to metabolic reprogramming and this causes downstream Wnt activation.

The preserved mitochondrial mass, membrane potential, and maximal respiratory capacity in BOS fibroblasts indicate a regulated metabolic reprogramming rather than organelle dysfunction, distinguishing BOS from primary mitochondrial diseases. Notably, stable isotope tracing demonstrated maintained incorporation of glucose-derived carbon into TCA cycle intermediates despite reduced MPC abundance, suggesting compensatory adaptation through increased glycolytic throughput that sustains mitochondrial carbon supply. This compensatory model is supported by elevated aspartate, glutamate, acetyl-carnitine, and pentose phosphate pathway metabolites, reflecting broad metabolic rewiring beyond glycolysis alone.

Mechanistically, truncated ASXL1 and BAP1 show aberrant co-occupancy at an H3K4me3-marked intronic regulatory element within *MPC2* intron 1, characterized by broadened ASXL1 occupancy extending beyond BRD4-defined active regulatory boundaries while BRD4 remains focused and unchanged. This dissociation indicates loss of positional targeting specificity rather than altered transcriptional elongation complex activity. The canonical *MPC2* promoter double-peak ATAC-seq signature is preserved in BOS fibroblasts while the intronic element shows subtle increased accessibility, defining a two-component regulatory architecture in which chromatin perturbation at the intronic element influences transcript processing or alternative TSS usage rather than suppressing transcription initiation. The disproportionate protein-level reduction relative to modest transcript changes indicates additional post-transcriptional amplification through impaired heterodimer assembly, altered protein stability, or proteasomal turnover. Similar patterns of protein-level MPC suppression without overt transcriptional changes have been observed during late-gestation hypoxia, in hypertrophic hearts, and in endothelial cells under hypoxic stress ^49–51^, consistent with post-transcriptional amplification as a conserved mechanism of MPC regulation across distinct biological contexts.

### MPC loss recapitulates developmental and disease-associated metabolic states

Reduced MPC expression is observed across diverse biological contexts characterized by preferential glycolysis, including intestinal stem cell niches and hair follicle activation^19,52,21^. MPC loss in intestinal stem cells actively drives a less differentiated state by promoting proliferation and blocking differentiation, while iPSC reprogramming requires MPC downregulation for successful nuclear reprogramming. The absence of MPC reduction in BOS iPSCs and NPCs combined with its presence in differentiated BOS fibroblasts suggests that truncating *ASXL1* mutations impair the normal MPC upregulation accompanying differentiation, consistent with the established role of ASXL1 in maintaining cell-type-specific gene expression through polycomb regulation.

The clinical parallels are notable. MPC deficiency due to pathogenic *MPC1* variants (OMIM #614741) causes developmental delay, microcephaly, growth failure, and elevated serum lactate and pyruvate^53,54^. Some of these phenotypes, such as developmental delay and growth failure, overlap with BOS. However, serum pyruvate and lactate remained within normal limits, even during times of stress suggesting that the downregulation of MPC is less than that in MPC deficiency, allowing BOS patients to maintain at least some organismal homeostasis of pyruvate and lactate levels. The convergence of clinical and metabolic features supports functional significance of MPC dysregulation in BOS pathophysiology, with partial rather than complete MPC loss consistent with viability of affected individuals given that complete MPC1 knockout is embryonically lethal^23–25^. Particularly relevant is the BOS hair phenotype: pharmacologic or genetic MPC suppression activates hair follicle stem cell cycling and promotes hair growth^40,55^, providing a direct mechanistic explanation for the thick, rapidly growing hair that is among the most consistent clinical features of BOS.

While the mechanisms differ; germline MPC1 loss versus ASXL1 truncation-dependent MPC reduction, the convergence of clinical and metabolic features across these two disorders supports the functional significance of MPC dysregulation in BOS pathophysiology. Even without changes in MPC gene expression, altered pyruvate handling at the mitochondrial membrane can drive large-scale rewiring of metabolic substrate use, including increased reliance on pathways such as glutamate oxidation ^56^.

### Mitochondrial pyruvate restriction is upstream of Wnt signaling activation and nutrient sensing

Pharmacologic MPC inhibition induced AXIN2 expression and altered β-catenin in control cells, while direct Wnt activation increased glycolytic flux without suppressing mitochondrial respiration or reducing MPC levels, establishing that Wnt activation is a downstream metabolic stress response rather than the primary epigenetic output of ASXL1 truncation. This is consistent with literature demonstrating that mitochondrial dysfunction activates β-catenin signaling through altered acetyl-CoA availability^57^, cytosolic NAD⁺/NADH ratios^58^, and lactate-mediated signaling^59^. Beyond Wnt, BOS fibroblasts exhibited impaired mTOR-dependent nutrient sensing with attenuated pS6K under amino acid-replete and deprived conditions, partially rescued by L-asparagine, consistent with asparagine coupling mitochondrial respiration to mTOR–ATF4 signaling^45^. The absence of *ATF4* or *ASNS* transcriptional dysregulation confirms this impairment arises downstream of metabolic signaling rather than epigenetic silencing.

### PR-DUB complex hyperactivation converges on MPC dysregulation

Our findings extend the metabolic consequences of PR-DUB perturbation beyond what has been described for BAP1. Germline *BAP1* mutations drive a Warburg-like metabolic state in primary fibroblasts ^17,33^, and our data identify MPC loss as a candidate shared effector: ASXL1 and BAP1 co-occupancy at the *MPC2* intronic regulatory element provides a mechanistic basis for convergence, suggesting PR-DUB hyperactivity at this element, regardless of which subunit is perturbed, alters chromatin state in a way that reduces MPC2 protein output. Whether *BAP1* mutations similarly reduce MPC protein levels is a directly testable prediction that would implicate MPC dysregulation as a shared consequence of PR-DUB complex perturbation.

### Limitations and Future Directions

Our study provides the first characterization of metabolic reprogramming in patient-derived cells harboring germline *ASXL1* truncating variants, with several limitations informing interpretation. First, the cohort of seven BOS individuals reflects the extreme rarity of this disorder, with orthogonal validation in HEK293T and Caco-2 systems substantially mitigating this constraint. Second, the phenotype is differentiation-state dependent — absent in BOS iPSCs and NPCs but present in fibroblasts — suggesting ASXL1 truncation impairs normal MPC upregulation during differentiation. Third, while ChIP-seq and ATAC-seq define a two-component regulatory architecture at *MPC2* with aberrant intronic ASXL1/BAP1 occupancy dissociated from stable BRD4 positioning, the precise mechanism connecting this chromatin perturbation to reduced protein output requires isoform-resolved long-read RNA-seq, ribosome footprinting, or isoform-specific RT-PCR. Fourth, *in vivo* validation is absent; future studies in neural and cardiac model systems will determine how MPC dysregulation contributes to BOS manifestations including seizures, and whether similar mechanisms extend to *ASXL2*- and *ASXL3*-associated disorders.

Together, our findings establish a previously unrecognized pathway through which gain-of-function *ASXL1* truncating variants rewire cellular bioenergetics. Truncated ASXL1 and BAP1 accumulate aberrantly at an H3K4me3-marked intronic regulatory element within *MPC2*, reducing mitochondrial pyruvate carrier protein levels and driving a Warburg-like metabolic state characterized by increased glycolytic flux and accumulation of pyruvate and lactate. Pharmacologic MPC inhibition phenocopies the metabolic and Wnt signaling changes in ASXL1-mutant cells, while Wnt activation alone is insufficient to recapitulate the full metabolic phenotype, placing mitochondrial pyruvate restriction causally upstream of signaling dysregulation. These findings expand the functional repertoire of ASXL1 beyond transcriptional regulation, identify MPC as a critical effector linking epigenetic perturbation to metabolic–signaling crosstalk, and establish mitochondrial pyruvate import as a metabolic checkpoint through which chromatin-associated mutations can fundamentally alter cellular identity, with implications for therapeutic targeting in both Bohring–Opitz syndrome and *ASXL1*-mutant myeloid malignancy.

## METHODS

### Human fibroblast samples

Primary dermal fibroblasts were collected as outlined in Lin et al. (2023)^5^, which provides comprehensive details on the selection and characterization of cell lines under IRB-approved protocols with informed consent. To ensure representative diversity, we incorporated both male and female patients and controls, to minimize the potential for sex-chromosome-specific variations in our analysis (**Table 1**). Furthermore, we included cell lines derived from patients with different genetic variants, all of which have been classified as ‘likely pathogenic’ or ‘pathogenic’ by clinical laboratorians and have clinical features consistent with BOS.

### Cell culture and media conditions

Fibroblasts were maintained under typical growth conditions, as previously described ^5^. Cells were maintained in DMEM supplemented with 10% FBS, 1% penicillin-streptomycin and 1% non-essential amino acids (NEAA) unless otherwise specified. Catalogue numbers for each reagent are listed (**Supplemental Table 17**). Human embryonic kidney (HEK293T) and colorectal adenocarcinoma (Caco-2) were maintained in DMEM with 10% FBS and 1% penicillin-streptomycin (**Supplemental Table 1**). All cell lines were confirmed mycoplasma-free and used within 10 passages.

### Generation of EGFP-tagged mutant and full length ASXL1 plasmids

We developed EGFP tagged ASXL1 constructs in a pDest vector backbone in order to determine the effect of overexpression of ASXL1 on the cells, where expression is under a CMV promoter. We used pENTR1A backbone and built entry vectors using NEBuilder HiFi DNA Assembly (NEBuilder® HiFi DNA Assembly Cloning Kit, Catalog #E5520S, New England Biolabs) that contain the FKBP12(F36V) degron (from Addgene Plasmid #154287) with EGFP (from Addgene Plasmid #185763) and a flexible linker between EGFP and ASXL1 to allow for ASXL1 to be tagged at the N-term. Gateway Cloning (Invitrogen™ Gateway™ LR Clonase™ II Enzyme mix, Catalog #11-791-020, FisherScientific) was performed to develop the final pDest destination vectors. Plasmid containing the cDNA for truncated ASXL1 with 1-479 amino acids was built from Addgene Plasmid #74262, while cDNA for truncated ASXL1 with 1-1304 amino acids was built from Addgene Plasmid #74244. The full length ASXL1 cDNA was built from Addgene Plasmid #74244 and an Integrated DNA Technologies gBlock for 1305-1541 amino acids of ASXL1. A plasmid with three stop codons inserted at the end of the EGFP sequence and before the linker and full-length ASXL1 sequence was developed as a size control. The length of this construct is comparable to the full-length plasmid, but produces a product that is equivalent to the FKBP12(F36V) degron and EGFP only vector.

### Transfection of mutant ASXL1 plasmids and ASXL1 siRNAs

Using Lipofectamine™ 3000, we transfected the following cell types: HEK293T:

- Empty vector controls:

- 2.5ug of EGFP control

▪ pCMV-eGFP-NMHC- IIAdeltaIQ2 (Addgene, Plasmid #35690)
- Truncated ASXL1 constructs:

- 2.5µg of EGFP-tagged ASXL 1-479 (denoted ASXL1^GFP-1–479^)
- 2.5µg of EGFP-tagged ASXL 1-1304 (denoted ASXL1^GFP-1–1304^)
- Full length ASXL1:

- 2.5µg of EGFP-tagged ASXL 1-1541 (denoted ASXL1^GFP-1–1541^) Caco-2:
- Empty vector controls:

- 2.5ug of EGFP control

▪ pCMV-eGFP-NMHC- IIAdeltaIQ2 (Addgene, Plasmid #35690)
- Truncated ASXL1 constructs:

- 2.5µg of FLAG-tagged ASXL 1-479 (denoted ASXL1^1–479^)

▪ Addgene, Plasmid #74262
- 2.5µg of FLAG-tagged ASXL 1-1304 (denoted ASXL1^1–1304^)

▪ Addgene, Plasmid #74244

Transfected cells were incubated for 48 hours and transfection was confirmed with imaging and western blots **(Supplemental Figures 3, 4, and 13, Supplemental Tables 18 - 19)**. We then performed RNA-seq to validate transfection and identify differentially expressed genes.

Chemical inhibitors and treatments:

To modulate metabolic and signaling pathways, we used:

- MPC inhibition: UK5099 (Selleckchem Cat #S5317) at 1.25 µM, 2.5 µM, and 5 µM for 24 hours
- Wnt activation: CHIR99021 (Tocris Bioscience Cat #4423) at 2.5 µM, 5 µM, and 10 µM for 24 hours
- 0.05% DMSO as control

### Western blotting and protein quantification

Whole cell lysates were prepared as previously described ^5^, using a 10X Cell Lysis Buffer (Cell Signaling Technology, Catalog #9803S) supplemented with Halt Protease and Phosphatase Inhibitor Cocktail (100X, Thermo Fisher Scientific, Catalog #78440). Lysates were quantified using the Pierce BCA Protein Assay Kit (Thermo Fisher Scientific 23225).

Western blotting was performed using standard protocols ^60^. The commercial antibodies used are detailed in **Supplemental Table 18**. Uncropped blots are provided in **Supplemental Data**.

Protein bands were quantified using *ImageJ* software, facilitating the analysis of protein intensity relative to intensity of control protein, β-Actin, or tubulin. Data analysis was performed in RStudio, where protein intensity values were normalized against actin intensity values. Statistical evaluations, including pairwise t-tests, were employed to discern significant differences between the experimental conditions. Visualization of the results was performed using *ggplot2* in R. Normalized protein intensity was calculated relative to loading control. Boxplots depict normalized protein intensity fold change of target protein levels compared to control samples.

### Seahorse assay

Fibroblast cell lines were cultured under standard conditions and were seeded at 10,000 or 12,000 cells per well in an Agilent Seahorse 96 well XFe96 / XF Pro Microplate (Agilent 103794-100) one day prior to analysis. Each sample was seeded in at least triplicates. HEK293T cells were transfected with the EGFP-tagged mutant and full length ASXL1 plasmids. GFP positive cells were then sorted 24 hours post-transfection and reseeded at 35,000 cells per well in an Agilent Seahorse 96 well XFe96 / XF Pro Microplate (Agilent 103794-100) then analyzed 24 hours later.

Mitochondrial respiration and glycolysis were measured using an Agilent Seahorse XF96 Analyzer. The Seahorse XF Analyzer was loaded with pre-calibrated sensor cartridges containing compounds targeting mitochondrial respiration: oligomycin, FCCP, and a mixture of rotenone and antimycin A **(Supplemental Table 1)**. The Seahorse assay was conducted according to the manufacturer’s guidelines in Seahorse media containing 5 mM glucose, 2 mM glutamine, and 1 mM pyruvate, measuring oxygen consumption rate (OCR) and extracellular acidification rate (ECAR) at basal conditions, followed by sequential injections of oligomycin ( 2uM), FCCP (2 uM), and rotenone (1 uM) and antimycin A (2 uM).

Data were normalized to cell number per well by imaging nuclei with 5 ug/mL Hoechst. Data were analyzed using Wave software and visualized with Prism. Full trace data are provided in **Supplemental Data**.

### Mitochondrial imaging

To examine mitochondrial dynamics, we plated 12,000 fibroblast cells per well in a 96 well plate (Corning Costar 3603). Each sample was seeded in at least triplicates. We used tetramethylrhodamine, ethyl ester (TMRE) to assess mitochondrial membrane potential, and Mitotracker Green (MTG) to estimate mitochondrial mass. Initially, we added a carbonyl cyanide-4 (trifluoromethoxy) phenylhydrazone (FCCP) control to each well to induce depolarization of the mitochondrial membrane, serving as a crucial experimental control. Changes in mitochondrial membrane potential were quantified by measuring the fluorescence intensity of TMRE before and after FCCP treatment. This approach allowed us to analyze FCCP-induced depolarization effectively, providing insights into mitochondrial functionality under stress conditions.

### RNA sequencing (RNA-seq)

Fibroblasts were cultured as described above. Cells were harvested, RNA was extracted, and RNA-seq was conducted and analyzed according to established protocol ^5^. RNA-seq data are deposited in the NCBI Gene Expression Omnibus database: GSE230686 (fibroblast RNA-seq) and GSE337675 (Caco-2 and HEK293T RNA-seq).

### Intracellular metabolite extraction and analysis

Fibroblast cells were seeded in six-well plates at 200,000 cells per well, with at least three replicates per sample, per time point. At the following timepoints prior to extraction - 6 hours, 24 hours, 72 hours - media was changed and glucose tracing with [U-^13^C_6_]-glucose or glutamine tracing [U-^13^C_6_]-glutamine media were added. Metabolites were extracted at 70–80% confluence per established protocol ^45^. Briefly, after two washes with ice-cold 150 mM ammonium acetate, pH 7.3, 500 uL 80% methanol was added to each well. After incubation for 20 minutes at −80°C, cells were scraped off the plate, vortexed vigorously, and centrifuged at maximum speed. 250 uL of the resulting supernatant was dried under vacuum. Dried metabolites were stored at -80°C. Mass spectrometry analysis was conducted per established protocol ^45^ and samples were run on a Vanquish (Thermo Scientific) UHPLC system coupled to a Q-Exactive (Thermo Scientific) mass analyzer.

For stable isotope tracing experiments, the peak areas were additionally processed via the R package AccuCor ^61^ to correct for natural isotope abundance. Peak areas for each sample were normalized by the measured area of the internal standard trifluoromethane- sulfonate (present in the extraction buffer) and by the number of cells present in the extracted well. Analysis was conducted using the Christofk Lab Grapha platform. Significantly altered metabolites were identified using p adjusted < 0.05.

### Quantification and statistical analysis

Each experimental condition was repeated at least three times. Fibroblast lines from controls and BOS patients were seeded in technical replicates of at least three in each experimental condition. All experiments were performed in biological triplicate unless otherwise indicated.

Graphs present the mean +/-SD, and p values were generated by two-tailed Student’s t test. Asterisks indicate the significance of the p value: ∗p < 0.05; ∗∗p < 0.01; ∗∗∗p < 0.001. Data were analyzed and visualized using GraphPad Prism and *ggplot2*.

## Data availability

All metabolomic, Seahorse and western blot quantification data are provided in the Supplemental figures, tables and data. Raw data is available upon request. The datasets generated during this study are available at the Gene Expression Omnibus (GEO) under the accession number GSE337675. Publicly available RNA-seq and ATAC-seq data from our prior study were reanalyzed: GSE230686 (RNAseq) and GSE230688 (ATAC-seq). The following public datasets for ChIP-seq were downloaded and reanalyzed: GSE302969 (ASXL1, BAP1, and BRD4), ENCFF439DDQ (H3K4me3) ENCFF885SUR (H3K27ac). This paper does not report original code. Any additional information required to reanalyze the data reported in this paper is available from the lead contact upon request.

## ACKNOWLEDGEMENTS

We thank Dr. Cristian Beninca for respirometry, extracellular flux analysis and mitochondrial imaging. We acknowledge the contributions for the UCLA California Center for Rare Disease and Dr. Stanley Nelson for his support. We also thank all the patients and their families for their support and contributions to our research.

## CONTRIBUTIONS

**[IL]:** Conceptualization, Data Generation, Formal Analysis, Writing-original draft. **[MR]:** Data Generation, Formal Analysis, Writing-original draft, **[AK]:** Data Generation and Formal Analysis, **[NV]:** Resources, **[SS]:** Data Curation, **[NM]:** Methodology, **[AN]:** Data Generation, **[LS]:** Methodology, **[BER]:** Supervision, Writing-review and Editing, **[PMBM]:** Supervision, **[HC]:** Conceptualization, Resources, Methodology, **[VAA]:** Conceptualization, Supervision, Funding acquisition, Writing- review and editing.

