## Supplemental Figure Legends for "Truncating *ASXL1* variants rewire cellular metabolism via mitochondrial pyruvate carrier repression"

**Supplemental Figure 1. BOS fibroblasts exhibit increased glycolytic ATP production and reduced relative contribution of oxidative phosphorylation**

Seahorse ATP rate analysis was performed in patient-derived BOS fibroblasts and unaffected control fibroblasts to assess the relative contribution of glycolysis and mitochondrial oxidative phosphorylation to cellular ATP production.
**(A)** Glycolysis-derived ATP production (ATP glycolysis) is significantly increased in BOS fibroblasts compared with controls.
**(B)** Absolute mitochondrial ATP production (ATP OXPHOS) is not significantly different between BOS and control fibroblasts.
**(C)** The proportional contribution of mitochondrial ATP production to total cellular ATP is significantly reduced in BOS fibroblasts, indicating a relative shift toward glycolytic ATP generation.
Data are shown as mean ± s.e.m. Statistical significance was assessed using one-way ANOVA with post hoc multiple-comparison correction.

**Supplemental Figure 2. ASXL1 truncation does not overtly alter mitochondrial abundance or morphology in BOS fibroblasts**

Representative fluorescence microscopy images of control and BOS patient-derived fibroblasts stained with MitoTracker Green to assess mitochondrial mass and tetramethylrhodamine ethyl ester (TMRE) to evaluate mitochondrial membrane potential. Patient identifiers are indicated. No overt differences in mitochondrial abundance, intracellular distribution, or mitochondrial morphology were observed between control and BOS fibroblasts under the conditions examined. Scale bars, 500 μm.

**Supplemental Figure 3. Validation of HEK293T transfection efficiency and ASXL1 construct expression**

To validate ASXL1 expression constructs and transfection efficiency, HEK293T cells were transiently transfected with GFP-tagged control and ASXL1 expression plasmids.

**(A)** Schematic representation of the expression constructs used in this study, including GFP control, ASXL1 1–479, ASXL1 1–1304, and full-length ASXL1 1–1541 plasmids. Predicted protein sizes and construct architecture are indicated.

**(B)** Representative brightfield and fluorescence microscopy images of fluorescence-activated cell sorted (FACS) GFP-positive HEK293T cells following transfection at 20X. GFP fluorescence confirms successful enrichment of transfected cells prior to downstream metabolic and biochemical analyses.

Scale bars as indicated.

Transfection efficiencies

Plasmid size control (GFP): 60.7%

1-479: 29.4%

1-1304: 20%

1-1541: 55%

**Supplemental Figure 4. Immunoblot validation of ASXL1 construct expression in HEK293T cells**

Western blot validation of ASXL1 construct expression in transfected HEK293T cells. Protein lysates from untransfected cells, mock-transfected controls, and cells expressing GFP-tagged ASXL1 constructs were probed for GFP, with β-actin used as a loading control. Multiple GFP-reactive bands were detected in cells expressing truncated and full-length ASXL1 constructs, with the predominant bands corresponding to the expected molecular weights of the respective ASXL1 protein products. These findings confirm successful expression of ASXL1 truncating and full-length constructs.

**Supplemental Figure 5. Aberrant ASXL1 expression promotes glycolytic ATP production and reduces the relative contribution of oxidative phosphorylation**

Seahorse ATP rate analysis was performed in HEK293T cells expressing full-length or truncating ASXL1 constructs and compared with empty-vector controls.
**(A)** Glycolysis-derived ATP production (ATP glycolysis) is significantly increased in ASXL1-expressing cells relative to controls.
**(B)** Absolute mitochondrial ATP production (ATP OXPHOS) is also increased following ASXL1 expression.
**(C)** The proportional contribution of oxidative phosphorylation to total ATP production is reduced in ASXL1-expressing cells, indicating a relative shift toward glycolytic ATP generation.
Data are shown as mean ± s.e.m. Statistical significance was assessed using one-way ANOVA with post hoc multiple-comparison correction.

**Supplemental Figure 6. Untargeted metabolomics identifies elevated glycolytic intermediates in BOS fibroblasts**

Untargeted metabolomic profiling was performed in BOS and control fibroblasts to identify metabolic alterations associated with ASXL1 truncation.
**(A)** Volcano plot showing differential metabolite abundance in BOS fibroblasts relative to controls. Significantly altered metabolites are highlighted. Significant metabolites defined by adjusted P < 0.05.
**(B)** Heatmap of glycolytic intermediates detected by untargeted metabolomics demonstrates altered metabolic profiles in BOS fibroblasts. Red dots indicate metabolites significantly altered between groups (P < 0.05).
**(C)** Relative abundance of intracellular glucose is significantly increased in BOS fibroblasts compared with controls.
**(D)** Relative abundance of pyruvate is significantly increased in BOS fibroblasts compared with controls.
**(E)** Relative abundance of lactate is increased but does not reach statistical significance.
Data are shown as mean ± s.e.m. Statistical significance was determined using two-tailed Student’s t-tests with multiple-testing correction where applicable.

**Supplemental Figure 7. Stable glucose and glutamine carbon utilization in BOS fibroblasts following 24-hour isotope tracing**

Stable isotope tracing was performed using U-¹³C-glucose and U-¹³C-glutamine to assess carbon utilization in BOS and control fibroblasts following 24-hour labeling.
**(A)** Fractional labeling of intracellular glucose pools demonstrates comparable glucose isotopologue distributions between BOS and control fibroblasts, with approximately 70% M+6 and 30% M+0 labeling in both groups.
**(B)** U-13C-glucose tracing of intracellular pyruvate demonstrates predominantly complete M+3 labeling in both BOS and control fibroblasts, with no significant differences in isotopologue distribution.
**(C)** Fractional contribution of glucose-derived and glutamine-derived carbon to intracellular lactate pools is comparable between BOS and control fibroblasts.
Data are shown as mean ± s.e.m. Statistical significance was assessed using two-tailed Student’s t-tests.

**Supplemental Figure 8. Early isotope tracing reveals enhanced glycolytic and tricarboxylic acid cycle metabolite abundance in BOS fibroblasts**

Targeted metabolomic analysis with U-¹³C-glucose tracing was performed following 6-hour labeling to characterize early metabolic flux changes in BOS fibroblasts.
**(A)** Heatmap of glycolytic intermediates demonstrates increased abundance of multiple glycolytic metabolites in BOS fibroblasts relative to controls.
**(B)** Heatmap of tricarboxylic acid (TCA) cycle intermediates demonstrates increased abundance of several TCA-associated metabolites in BOS fibroblasts.
**(C)** Pyruvate isotopologue analysis demonstrates predominant M+3 labeling in both control and BOS fibroblasts, with complete glucose-derived carbon incorporation and no significant differences in fractional contribution.
**(D)** Lactate isotopologue composition is similarly dominated by M+3 labeling in both groups; however, BOS fibroblasts exhibit a significantly increased fractional contribution of glucose-derived carbon to intracellular lactate pools.
**(E)** Acetyl-CoA isotopologue composition is predominantly unlabeled (M+0) in both control and BOS fibroblasts, with no significant differences in isotopologue distribution or fractional contribution.
Data are shown as mean ± s.e.m. Statistical significance was assessed using one-way ANOVA with post hoc multiple-comparison correction. Isotopologue distributions are presented as fractional contribution of labeled carbon species, and metabolite abundance is shown as normalized pool size. All significance tests were performed on normalized data with a one-way ANOVA. * p-value < 0.05, ** p-value <0.01, *** p-value < 0.001.

**Supplemental Figure 9. Prolonged isotope tracing demonstrates persistent pyruvate accumulation with limited alterations in downstream carbon partitioning**

Targeted metabolomic analysis with U-¹³C-glucose tracing was performed following 24-hour labeling to characterize metabolic flux in BOS and control fibroblasts.
**(A)** Heatmap of glycolytic intermediates demonstrates persistent metabolic alterations in BOS fibroblasts, although fewer metabolites remain significantly altered compared with earlier time points. Pyruvate abundance remains significantly elevated in BOS fibroblasts.
**(B)** Heatmap of tricarboxylic acid (TCA) cycle intermediates demonstrates modest differences between BOS and control fibroblasts at 24 hours.
**(C)** Pyruvate isotopologue analysis demonstrates predominant M+3 labeling in both BOS and control fibroblasts, with no significant differences in isotopologue composition or glucose-derived carbon contribution.
**(D)** Lactate isotopologue composition remains predominantly M+3 in both groups, with no significant differences in fractional carbon contribution.
**(E)** Acetyl-CoA isotopologue analysis demonstrates approximately equal representation of labeling in both BOS and control fibroblasts, without significant differences in isotopologue distribution or fractional contribution.
Data are shown as mean ± s.e.m. Statistical significance was assessed using one-way ANOVA with post hoc multiple-comparison correction. Isotopologue distributions are presented as fractional contribution of labeled carbon species, and metabolite abundance is shown as normalized pool size. All significance tests were performed on normalized data with a one-way ANOVA. * p-value < 0.05, ** p-value <0.01, *** p-value < 0.001.

**Supplemental Figure 10. Metabolic alterations in BOS fibroblasts persist following extended isotope tracing**

Targeted metabolomic analysis with U-¹³C-glucose tracing was performed following 72-hour labeling to assess sustained metabolic remodeling in BOS fibroblasts.
**(A)** Heatmap of glycolytic intermediates demonstrates persistent upregulation of glycolytic metabolism in BOS fibroblasts relative to controls.
**(B)** Heatmap of tricarboxylic acid (TCA) cycle intermediates similarly demonstrates sustained metabolic differences between BOS and control fibroblasts, although fewer metabolites reach statistical significance at this later time point.
**(C)** Pyruvate isotopologue analysis demonstrates complete M+3 labeling in both BOS and control fibroblasts without significant differences in glucose-derived carbon contribution.
**(D)** Lactate isotopologue composition remains predominantly M+3 (~85%) in both groups, with no significant differences in isotopologue distribution or fractional contribution.
**(E)** Acetyl-CoA isotopologue composition and glucose-derived carbon contribution remain comparable between BOS and control fibroblasts.
Data are shown as mean ± s.e.m. Statistical significance was assessed using one-way ANOVA with post hoc multiple-comparison correction. Isotopologue distributions are presented as fractional contribution of labeled carbon species, and metabolite abundance is shown as normalized pool size. All significance tests were performed on normalized data with a one-way ANOVA. * p-value < 0.05, ** p-value <0.01, *** p-value < 0.001.

**Supplemental Figure 11. BOS fibroblasts exhibit altered amino acid, redox, and mitochondrial shuttle metabolism following prolonged U-¹³C-glucose tracing**

Targeted metabolomic analysis was performed following 72-hour U-¹³C-glucose tracing to assess metabolic pathways beyond glycolysis and the tricarboxylic acid (TCA) cycle in BOS and control fibroblasts.

**(A)** Metabolite set enrichment analysis identifies significant alterations in pathways associated with glutamate metabolism, the malate–aspartate shuttle, carnitine synthesis, and the pentose phosphate pathway in BOS fibroblasts relative to controls.

**(B)** Aspartate isotopologue analysis demonstrates increased fractional contribution of glucose-derived carbon in BOS fibroblasts together with increased total metabolite abundance, consistent with enhanced carbon flux through pathways linked to mitochondrial pyruvate metabolism and the malate–aspartate shuttle.

**(C)** Glutamate isotopologue analysis demonstrates increased fractional contribution of glucose-derived carbon and increased total glutamate abundance in BOS fibroblasts, indicating altered anaplerotic and amino acid metabolic pathways downstream of glycolysis.

**(D)** Isotopologue distribution and metabolite pool size analysis of NAD⁺ demonstrates a significant increase in total NAD⁺ abundance in BOS fibroblasts, consistent with altered redox metabolism.

**(E)** Acetyl-carnitine isotopologue analysis demonstrates approximately equal proportions of M+2 and M+0 labeling in both BOS and control fibroblasts, consistent with mixed contributions from glucose-derived and unlabeled carbon sources. Total acetyl-carnitine abundance is increased in BOS fibroblasts.

**(F)** Sorbitol isotopologue analysis demonstrates predominant M+6 labeling in both BOS and control fibroblasts, indicating glucose-derived carbon incorporation through the polyol pathway. Total sorbitol abundance is significantly increased in BOS fibroblasts.

Data are shown as mean ± s.e.m. Statistical significance was assessed using one-way ANOVA with post hoc multiple-comparison correction. Isotopologue distributions are presented as fractional contribution of labeled carbon species, and metabolite abundance is shown as normalized pool size. All significance tests were performed on normalized data with a one-way ANOVA. * p-value < 0.05, ** p-value <0.01, *** p-value < 0.001.

**Supplemental Figure 12. ASXL1 truncation reduces MPC1 protein abundance in HEK293T cells**

Western blot analysis was performed in cytoplasmic lysates from HEK293T cells transfected with control plasmid or ASXL1 truncating constructs (ASXL1 1–479 and ASXL1 1–1304).
**(A)** Representative immunoblots of MPC1, MPC2, and asparagine synthetase (ASNS) in control and ASXL1-transfected cells shown in biological triplicate. ASNS abundance remained unchanged across conditions.
**(B)** Quantification demonstrates significantly reduced MPC1 protein abundance following expression of ASXL1 truncating constructs, whereas MPC2 abundance was not significantly altered.
Protein abundance was normalized to loading controls and expressed relative to control cells. Data are shown as mean ± s.e.m. Statistical significance was assessed using one-way ANOVA with post hoc multiple-comparison correction.

**Supplemental Figure 13. Validation of ASXL1 construct expression in CACO2 cells**

Validation of ASXL1 construct expression was performed in CACO2 cells transfected with GFP control or truncating ASXL1 constructs.
**(A)** Representative western blot analysis of nuclear protein extracts confirms expression of truncated ASXL1 proteins.
**(B)** Representative western blot analysis of cytoplasmic extracts demonstrates compartment-specific protein distribution following transfection.
**(C)** RNA sequencing-derived normalized ASXL1 transcript abundance demonstrates significant upregulation of ASXL1 expression in cells transfected with truncating ASXL1 constructs relative to controls.
Data are shown as mean ± s.e.m. Statistical significance was assessed using one-way ANOVA with post hoc multiple-comparison correction.

**Supplemental Figure 14. Independent biological replicate experiments confirm metabolic effects of MPC inhibition and Wnt activation**

Independent biological replicate experiments were performed to validate the effects of mitochondrial pyruvate carrier inhibition and Wnt pathway activation on cellular metabolism.

**(A–D)** Seahorse extracellular flux analysis following treatment with the mitochondrial pyruvate carrier inhibitor UK5099. Representative traces demonstrate **(A)** extracellular acidification rate (ECAR) and **(B)** oxygen consumption rate (OCR) across increasing UK5099 concentrations. Quantification of **(C)** basal ECAR demonstrates a dose-dependent increase in glycolytic activity, while **(D)** the basal OCR/ECAR ratio is significantly reduced with increasing UK5099 concentration, confirming a shift toward glycolytic metabolism.

**(E–H)** Seahorse extracellular flux analysis following treatment with the Wnt pathway agonist CHIR99021. Representative traces demonstrate **(E)** ECAR and **(F)** OCR across increasing CHIR99021 concentrations. Quantification of **(G)** basal ECAR demonstrates a dose-dependent increase in glycolytic activity, while **(H)** the basal OCR/ECAR ratio is significantly reduced with increasing CHIR99021 concentration, consistent with enhanced glycolytic metabolism.

Data are shown as box-and-whisker plots (median, interquartile range, and range) or mean ± s.e.m. ECAR is reported as mpH min⁻¹ and OCR as pmol O₂ min⁻¹. Statistical significance was assessed using one-way ANOVA with post hoc multiple-comparison correction. Exact P values are indicated in the corresponding panels.

**Supplemental Figure 15. BOS fibroblasts exhibit impaired mTOR signaling during amino acid stress**

Western blot analysis was performed to assess mTOR pathway activity in BOS and control fibroblasts following non-essential amino acid (NEAA) depletion and amino acid repletion.
**(A)** Following 16 hours of NEAA depletion, BOS fibroblasts exhibit significantly reduced phosphorylation of S6 kinase (pS6K) and reduced ATF4 abundance compared with controls under both amino acid-replete (+NEAA) and depleted (−NEAA) conditions.
**(B)** Following 48 hours of NEAA depletion, BOS fibroblasts exhibit further reduction in pS6K abundance relative to controls.
**(C)** L-asparagine repletion following 48 hours of NEAA depletion restores pS6K abundance in BOS fibroblasts.
**(D)** Following 72 hours of NEAA depletion, BOS fibroblasts demonstrate sustained suppression of mTOR signaling, particularly affecting the pS6K p85 isoform.
**(E)** Fold-change quantification of pS6K p70 abundance under NEAA depletion and L-asparagine repletion conditions corresponding to panels A–D.
**(F)** Fold-change quantification of pS6K p85 abundance under NEAA depletion and L-asparagine repletion conditions corresponding to panels A–D.
Protein abundance was normalized to loading controls and expressed relative to corresponding +NEAA conditions. Data are shown as mean ± s.e.m. Statistical significance was assessed using one-way ANOVA with post hoc multiple-comparison correction.

**Supplemental Figure 16. Altered nutrient signaling in BOS fibroblasts occurs despite preserved ATF4 and ASNS transcription**

**(A)** Representative CUT&RUN profiles for the active promoter mark H3K4me3 at the *ATF4* locus in control and BOS fibroblasts. BOS fibroblasts demonstrate increased H3K4me3 occupancy at the *ATF4* promoter relative to controls, indicating preserved or enhanced chromatin features associated with transcriptional activation.

**(B)** Differential chromatin accessibility analysis of an ATAC-seq peak located at the *ATF4* transcription termination site (TTS). BOS fibroblasts exhibit reduced accessibility at this region relative to controls (log₂FC = −0.87, adjusted P = 0.004).

**(C)** Normalized *ATF4* transcript abundance in BOS patient fibroblast RNA sequencing. No significant difference in *ATF4* expression was observed.

**(D)** Normalized *ASNS* transcript abundance in BOS patient fibroblast RNA sequencing. No significant differences in *ASNS* expression was observed.

**Supplemental Figure 17. β-catenin does not exhibit differential occupancy at MPC1 and MPC2 promoter regions following Wnt activation**

Publicly available β-catenin ChIP-seq datasets (GSE182842) were examined to determine whether canonical Wnt signaling directly regulates mitochondrial pyruvate carrier genes.

**(A)** β-catenin ChIP-seq tracks spanning the *MPC1* promoter region under basal (-Wnt) and Wnt-stimulated (+Wnt) conditions. Occupancy was comparable between conditions.

**(B)** β-catenin ChIP-seq tracks spanning the *MPC2* promoter region under basal (-Wnt) and Wnt-stimulated (+Wnt) conditions. Occupancy was comparable between conditions.

**Supplemental Figure 18. RNA sequencing quality control and transcriptomic validation in HEK293T ASXL1 models**

RNA sequencing quality control and global transcriptomic analyses were performed in HEK293T cells transfected with ASXL1 constructs and control plasmids.
**(A)** Dispersion estimates generated during DESeq2 analysis demonstrate appropriate mean–variance relationships across sequenced samples.
**(B)** Principal component analysis (PCA) of PC1 and PC2 demonstrates sample segregation primarily by sequencing batch (PC1, 78.85% variance), which was corrected during downstream DESeq2 analysis, and by transfection condition (PC2, 6.83% variance).
**(C)** PCA of PC3 and PC4 further resolves samples according to ASXL1 construct and transfection group, demonstrating reproducible transcriptional signatures associated with ASXL1 expression.
**(D)** Unsupervised hierarchical clustering and heatmap analysis demonstrate clear separation between control and ASXL1-transfected samples, with distinct clustering of truncating and full-length ASXL1 conditions.

**Supplemental Figure 19. ASXL1 constructs induce distinct transcriptional programs in HEK293T cells**

Differential gene expression and pathway enrichment analyses were performed in HEK293T cells expressing ASXL1 constructs.
**(A–C)** Volcano plots showing differential gene expression in HEK293T cells transfected with ASXL1 1–479 (**A**), ASXL1 1–1304 (**B**), and ASXL1 1–1541 (**C**) relative to controls. Significantly upregulated genes are shown in red (log2FC > 1.5, adjusted P < 0.05), while significantly downregulated genes are shown in blue (log2FC < −1.5, adjusted P < 0.05).
**(D–F)** Gene ontology enrichment analyses corresponding to each ASXL1 construct. ASXL1 1–479 (**D**) is enriched for neurogenesis- and neuronal development-associated pathways, ASXL1 1–1304 (**E**) demonstrates enrichment of telencephalon development programs, and ASXL1 1–1541 (**F**) demonstrates enrichment of proteasome-mediated and ubiquitin-dependent catabolic processes.

**Supplemental Figure 20. RNA sequencing quality control and transcriptomic validation in CACO2 ASXL1 models**

RNA sequencing quality control and transcriptomic analyses were performed in CACO2 cells expressing truncating ASXL1 constructs and corresponding controls.
**(A)** Dispersion estimates generated during DESeq2 analysis demonstrate appropriate variance modeling across sequenced samples.
**(B)** Principal component analysis (PCA) demonstrates robust segregation by transfection condition. PC1 (58% variance) separates mock-transfected cells from GFP controls and ASXL1-transfected cells, while PC2 (19% variance) further distinguishes truncating ASXL1-expressing samples from both control groups, indicating reproducible transcriptional effects associated with ASXL1 truncation.
**(C)** Unsupervised hierarchical clustering and heatmap analysis demonstrate clear separation of control and ASXL1-transfected samples.

**Supplemental Figure 21. Truncating ASXL1 variants alter mitochondrial and respiratory transcriptional programs in CACO2 cells**

Differential gene expression and pathway enrichment analyses were performed in CACO2 cells expressing truncating ASXL1 constructs.
 **(A)** Volcano plot demonstrating significantly altered genes in ASXL1-transfected cells relative to controls. Significantly dysregulated genes were defined using adjusted P < 0.05 and log2 fold-change thresholds as indicated.
 **(B)** Gene ontology enrichment analysis demonstrates significant dysregulation of pathways associated with mitochondrial respiration, respiratory chain complex assembly, and mitochondrial respirasome organization in ASXL1-expressing cells.

**Supplemental Figure 22. Integrated transcriptomic analysis reveals shared ASXL1-dependent transcriptional signatures across cell types**

Integrated RNA sequencing analysis was performed across HEK293T and CACO2 ASXL1 models to identify conserved transcriptional responses.
**(A)** Principal component analysis of integrated datasets demonstrates segregation by both cellular background and transfection condition. PC3 (0.2% variance) and PC4 (0.1% variance) further separate samples according to ASXL1 construct and control status independent of cell type.
**(B)** Unsupervised hierarchical clustering demonstrates separation by cell type while preserving distinct clustering according to plasmid transfection condition, indicating conserved ASXL1-associated transcriptional effects across models.

**(C)** Volcano plot showing differentially expressed genes identified from the integrated analysis. A total of 439 genes were significantly differentially expressed (adjusted P < 0.05), including 231 upregulated and 208 downregulated genes. Significantly upregulated genes meeting log2 fold change cutoff (log2FC > 0.58) are shown in red and significantly downregulated genes meeting log2 fold change cutoff (log2FC < -0.58) are shown in blue.

**(D)** Gene ontology enrichment analysis of significantly differentially expressed genes demonstrates enrichment of pathways related to protein–RNA complex assembly, cytoplasmic transport, and RNA transport.

**Supplemental Figure 23. Pyruvate dehydrogenase kinase family genes are not transcriptionally dysregulated following ASXL1 perturbation**

Expression of pyruvate dehydrogenase kinase (*PDK1–PDK4*) family genes was assessed across BOS fibroblasts and ASXL1 overexpression models to determine whether altered pyruvate metabolism was associated with transcriptional regulation of canonical pyruvate dehydrogenase inhibitory pathways.

**(A–D)** Normalized RNA sequencing transcript abundance of *PDK1*, *PDK2*, *PDK3*, and *PDK4* in BOS fibroblasts relative to control fibroblasts. No significant difference was observed.

**(E–H)** Normalized RNA sequencing transcript abundance of *PDK1*, *PDK2*, *PDK3*, and *PDK4* in CACO2 cells expressing ASXL1 truncation constructs relative to controls. No significant difference was observed.

**(I–L)** Normalized RNA sequencing transcript abundance of *PDK1*, *PDK2*, *PDK3*, and *PDK4* in HEK293T cells expressing ASXL1 constructs relative to controls. No significant difference was observed.

**Supplemental Figure 24. Glycolytic pathway genes are not transcriptionally dysregulated following ASXL1 perturbation**

Expression of glycolytic pathway genes were assessed across BOS fibroblasts and ASXL1 overexpression models to determine whether truncating ASXL1 mutations were associated with transcriptional dysregulation of glycolysis.

**(A–D)** Normalized RNA sequencing transcript abundance of key glycolytic pathway enzymes *HK1, HK2, PFKP and PKLR* in BOS fibroblasts relative to control fibroblasts. No significant difference was observed.

**(E–H)** Normalized RNA sequencing transcript abundance of key glycolytic pathway enzymes *HK1, HK2, PFKP* in CACO2 cells expressing ASXL1 truncation constructs relative to controls. No significant difference was observed.

**(I–L)** Normalized RNA sequencing transcript abundance of key glycolytic pathway enzymes *HK1, HK2, PFKP* in HEK293T cells expressing ASXL1 constructs relative to controls. No significant difference was observed.

**Supplemental Figure 25. Chromatin accessibility at MPC1 and MPC2 loci is preserved in BOS fibroblasts**

To determine whether reduced mitochondrial pyruvate carrier protein abundance in BOS fibroblasts is associated with altered chromatin accessibility, ATAC-seq profiles were examined at the *MPC1* and *MPC2* genomic loci in control and BOS patient-derived fibroblasts.

**(A)** Representative ATAC-seq tracks spanning the *MPC1* locus on chromosome 6. Three representative control fibroblast lines (n = 3 of 6 total controls) and three representative BOS fibroblast lines (n = 3 of 7 total BOS samples) are shown. Chromatin accessibility was assessed across both the 5′ (N-terminal) and 3′ (C-terminal) regions of the gene. Overall accessibility was comparable between groups, with a modest increase in accessibility observed within the C-terminal region of *MPC1* in BOS fibroblasts.

**(B)** Representative ATAC-seq tracks spanning the *MPC2* locus on chromosome 1. Three representative control fibroblast lines and three representative BOS fibroblast lines are shown. Prominent accessibility peaks were observed at the *MPC2* promoter region in both groups. BOS fibroblasts demonstrated more defined promoter-associated accessibility peaks relative to controls, consistent with preserved or enhanced chromatin accessibility at the *MPC2* locus.

Genomic coordinates and scale bars are indicated.
