## Supplemental Figures for "Truncating *ASXL1* variants rewire cellular metabolism via mitochondrial pyruvate carrier repression"

Supp Fig 1: BOS fibroblasts exhibit increased glycolytic ATP production and reduced relative contribution of oxidative phosphorylation

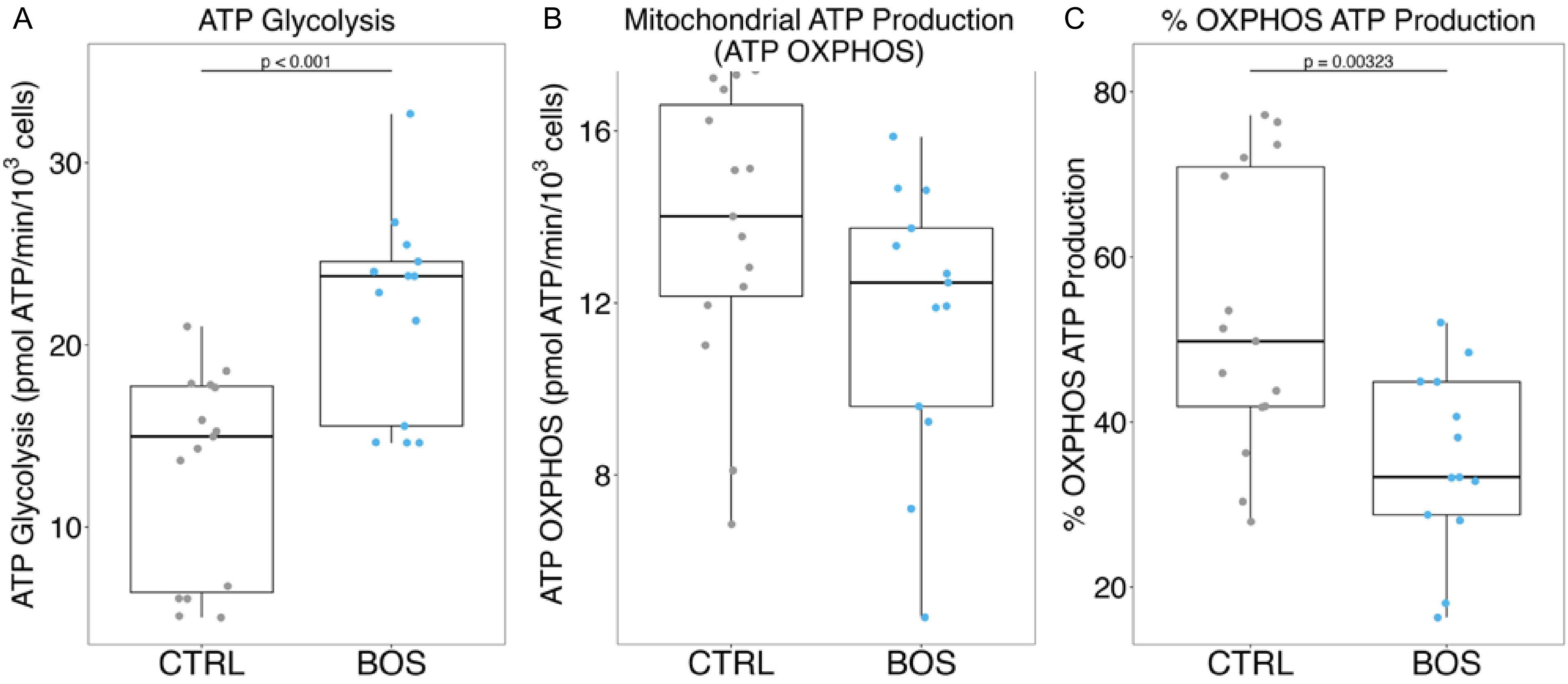

Supp Fig 2: ASXL1 truncation does not overtly alter mitochondrial abundance or morphology in BOS fibroblasts

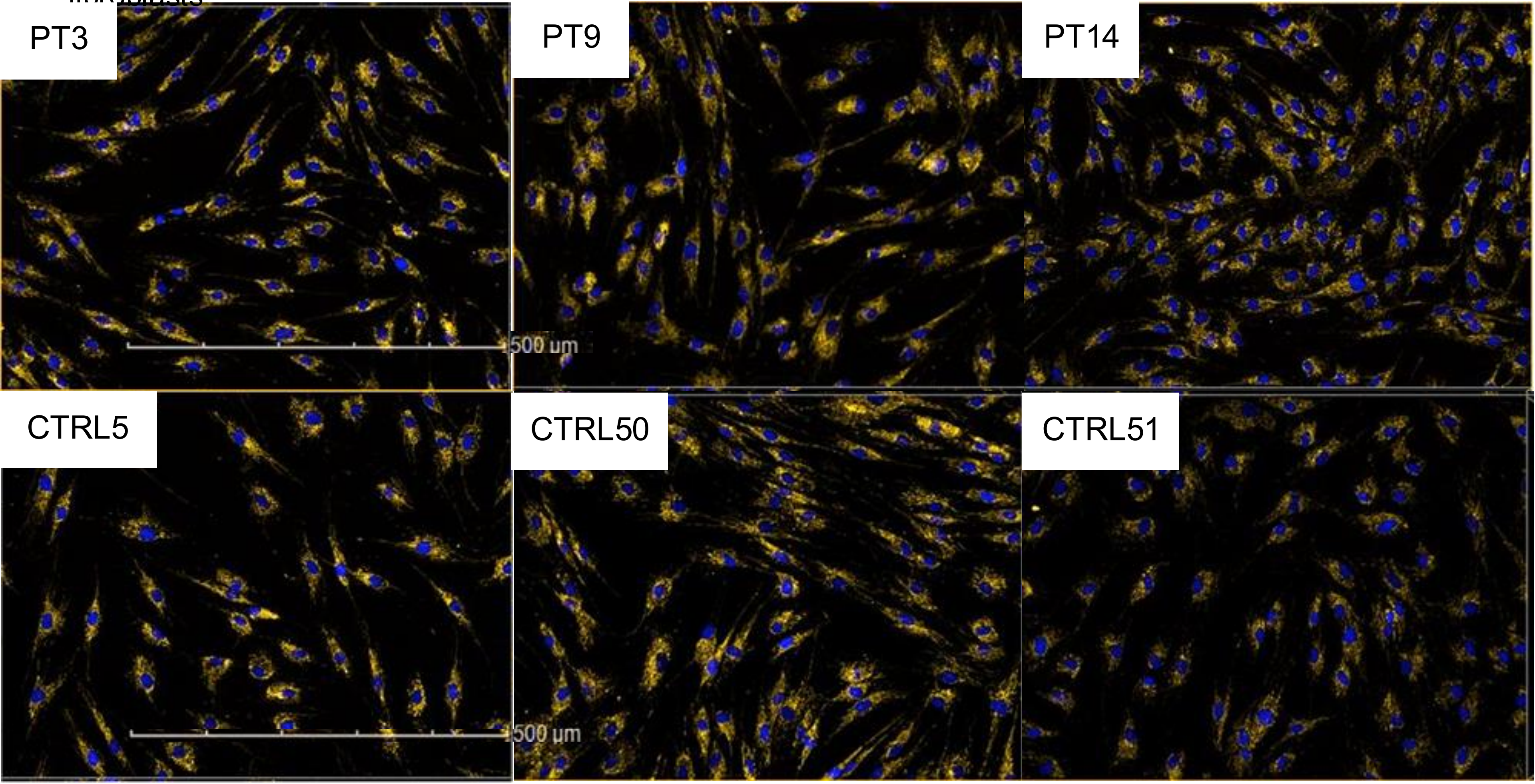

Supp Fig 3: Validation of HEK293T transfection efficiency and ASXL1 construct expression

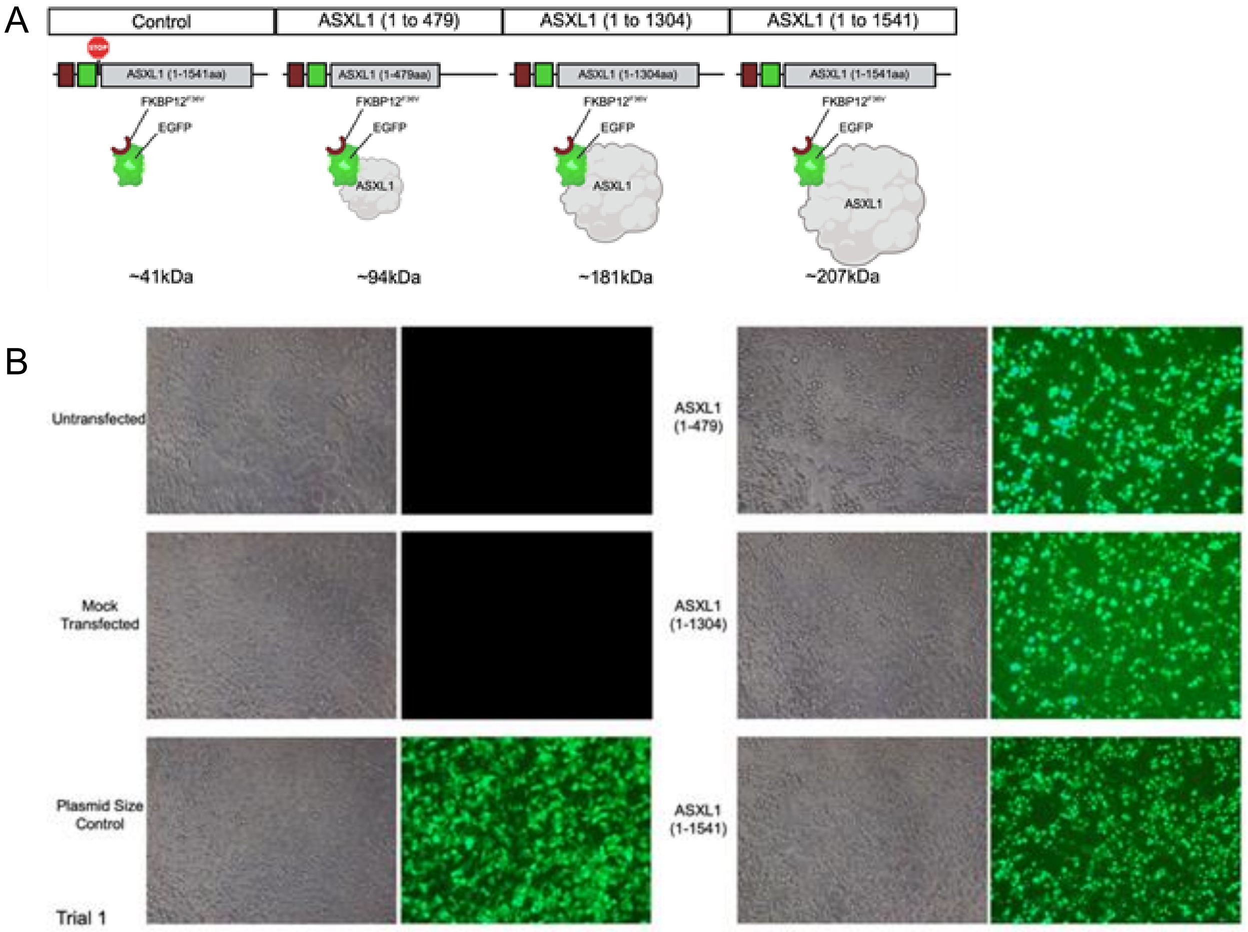

Supp Fig 4: Immunoblot validation of ASXL1 construct expression in HEK293T cells

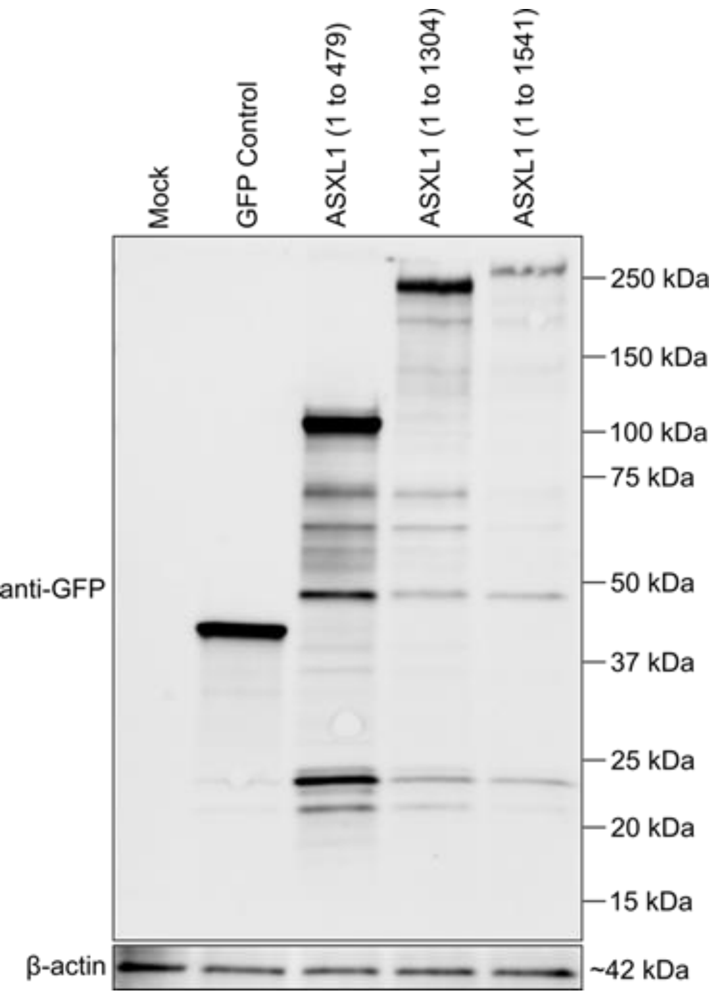

Supp Fig 5: Aberrant ASXL1 expression promotes glycolytic ATP production and reduces the relative contribution of oxidative phosphorylation

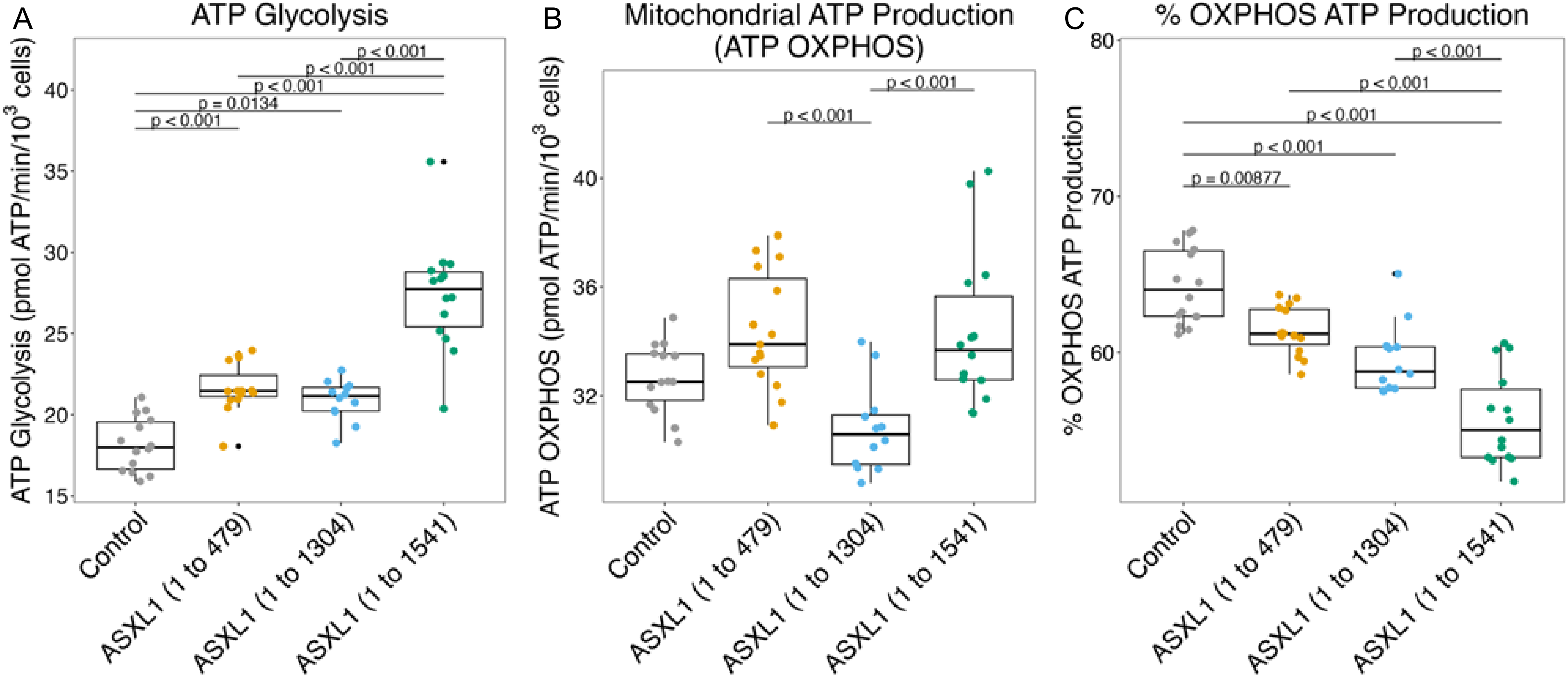

Supp Fig 6: Untargeted metabolomics identifies elevated glycolytic intermediates in BOS fibroblasts

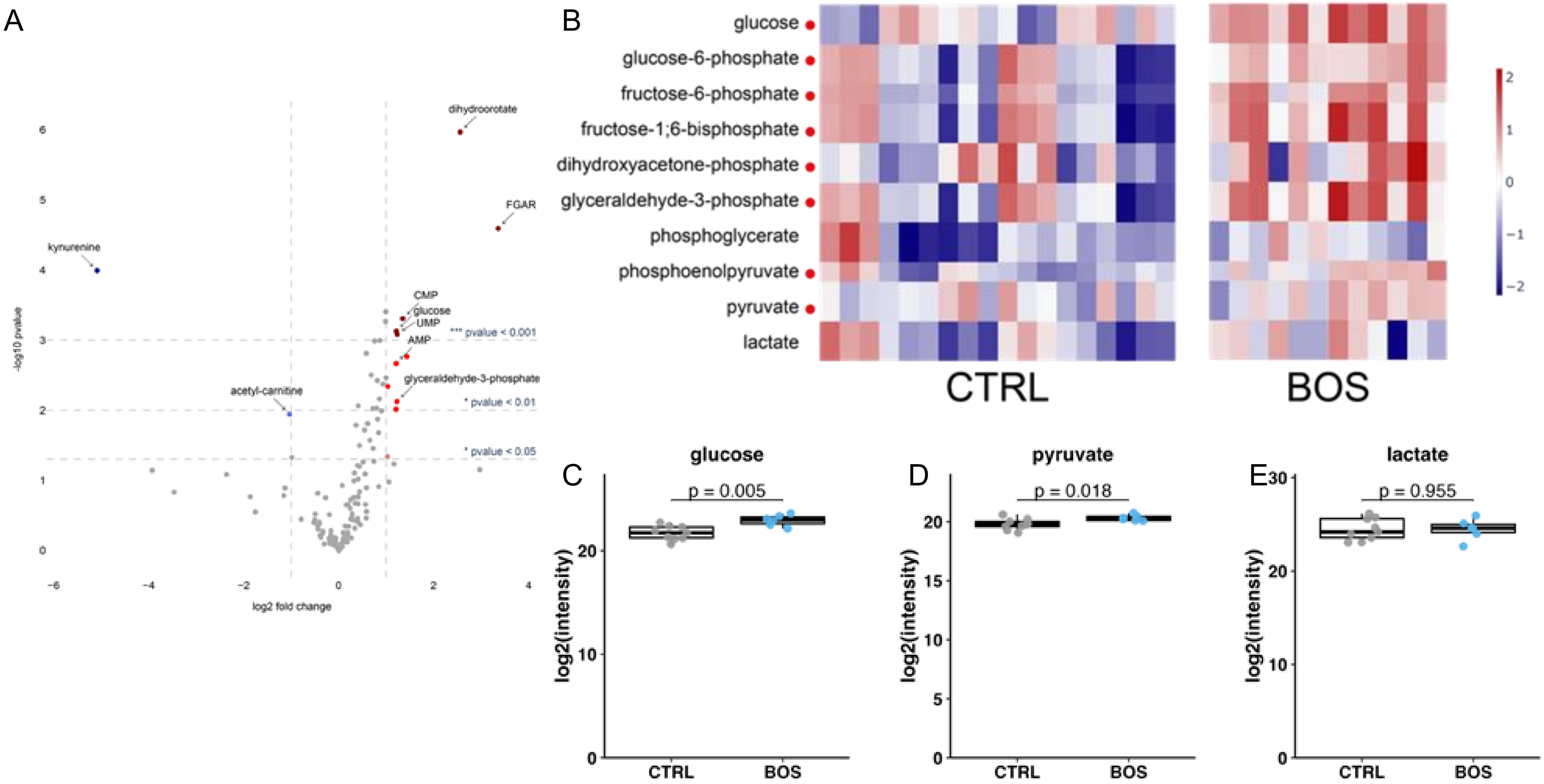

Supp Fig 7: Stable glucose and glutamine carbon utilization in BOS fibroblasts following 24-hour isotope tracing

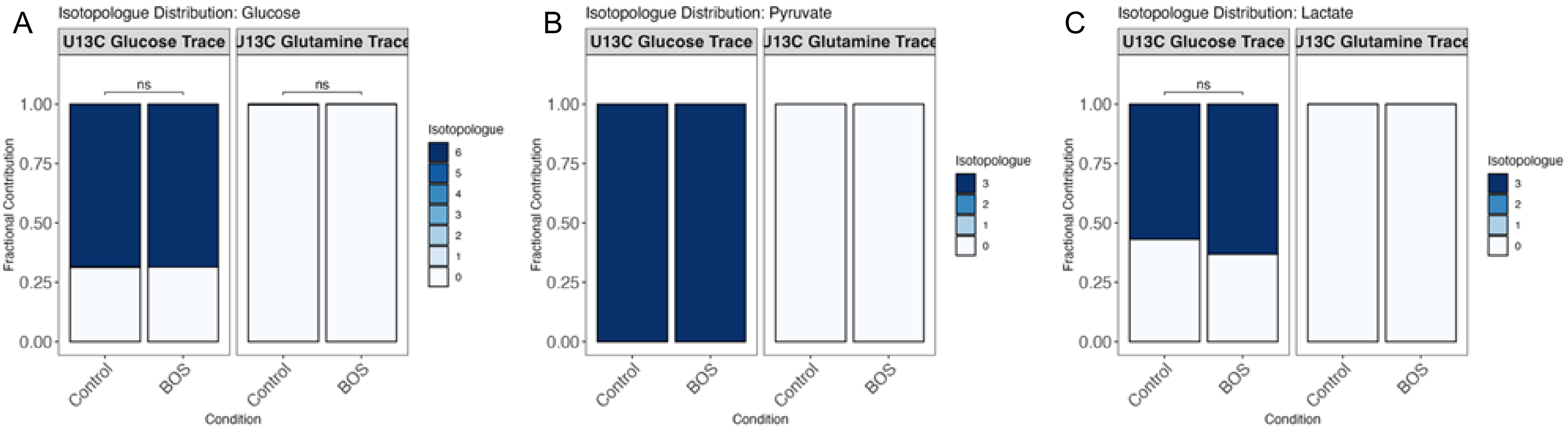

### Supp Fig 8: Early isotope tracing reveals enhanced glycolytic and tricarboxylic acid cycle metabolite abundance in BOS fibroblasts 6 hours after media change

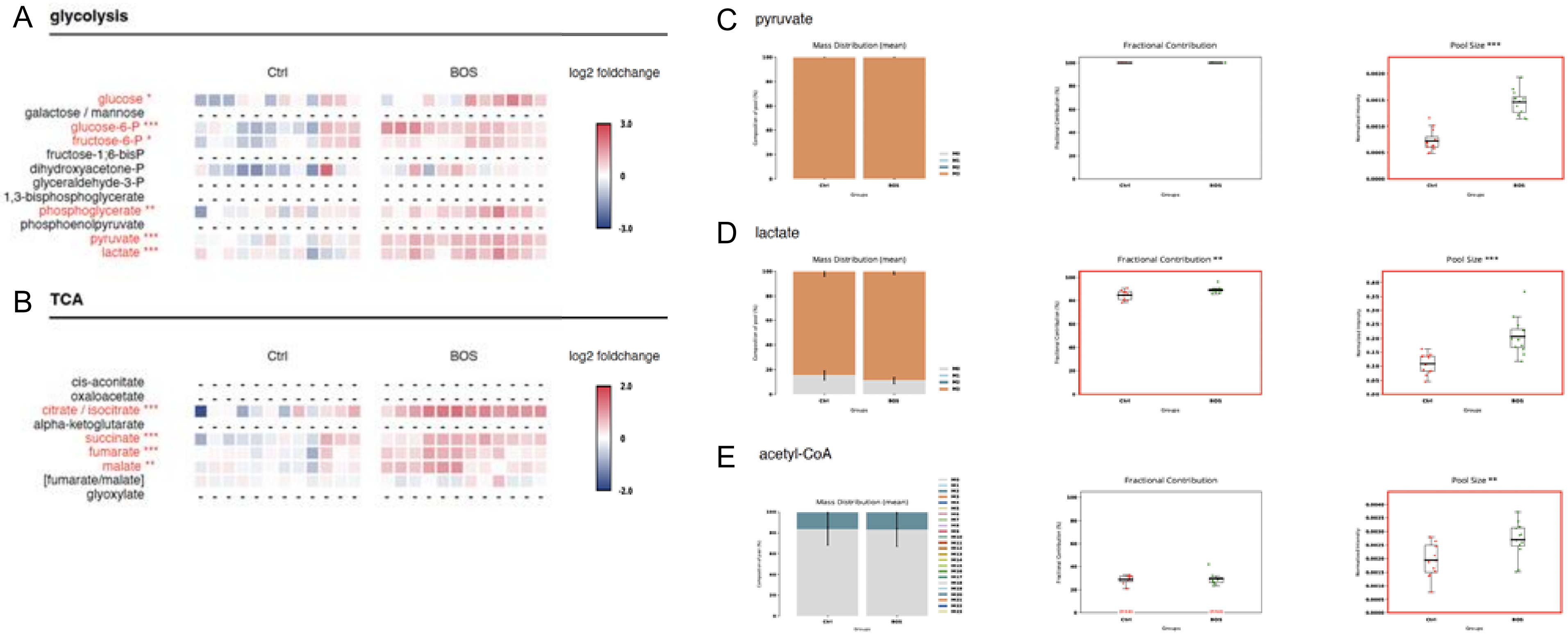

### Supp Fig 9: Prolonged isotope tracing demonstrates persistent pyruvate accumulation with limited alterations in downstream carbon partitioning 24 hours after media change

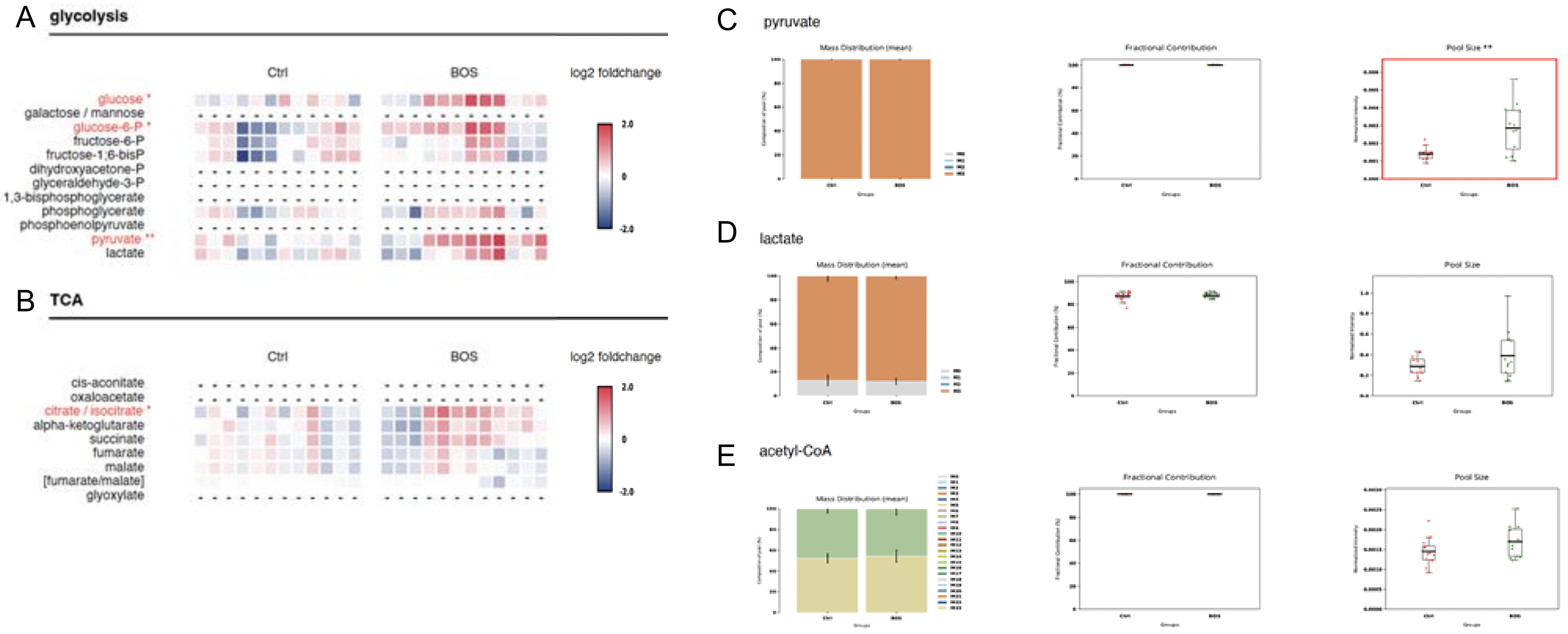

Supp Fig 10: Metabolic alterations in BOS fibroblasts persist following extended isotope tracing 72 hours after media change

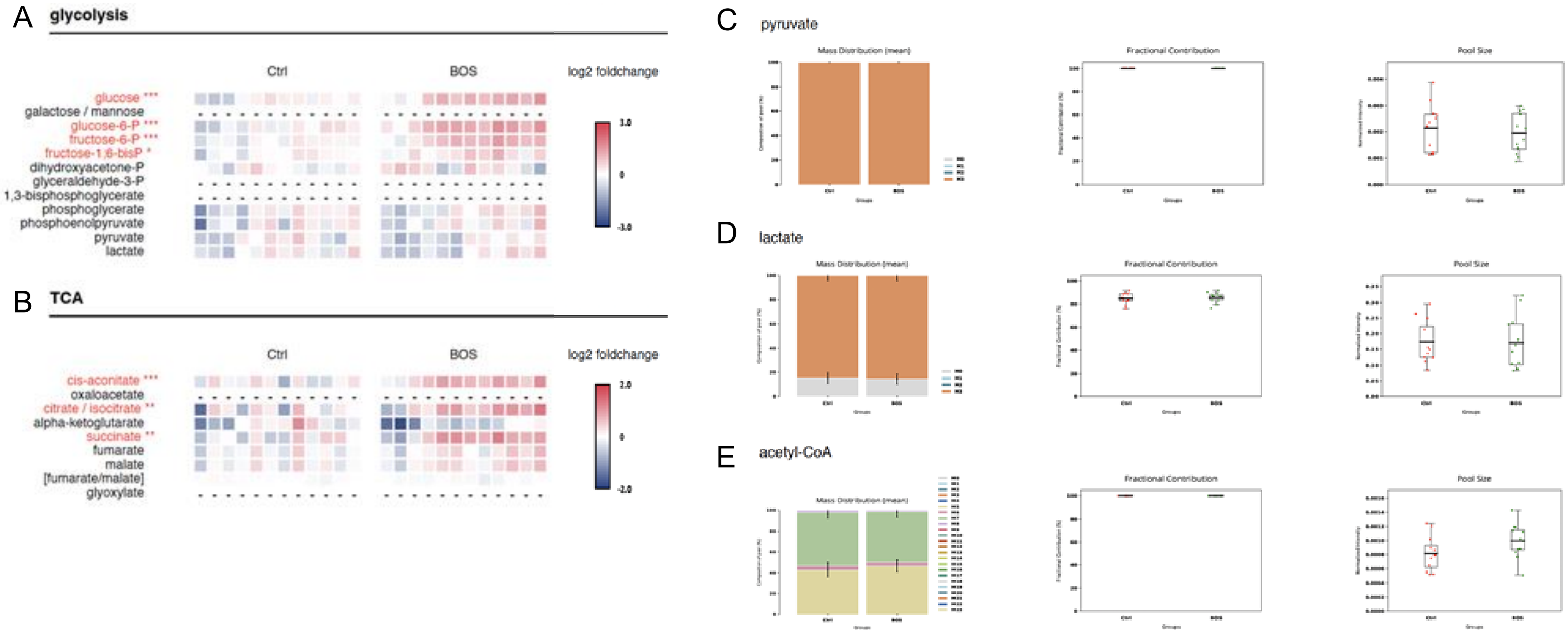

Supp Fig 11: BOS fibroblasts exhibit altered amino acid, redox, and mitochondrial shuttle metabolism following prolonged U-$^{13}$C-glucose tracing

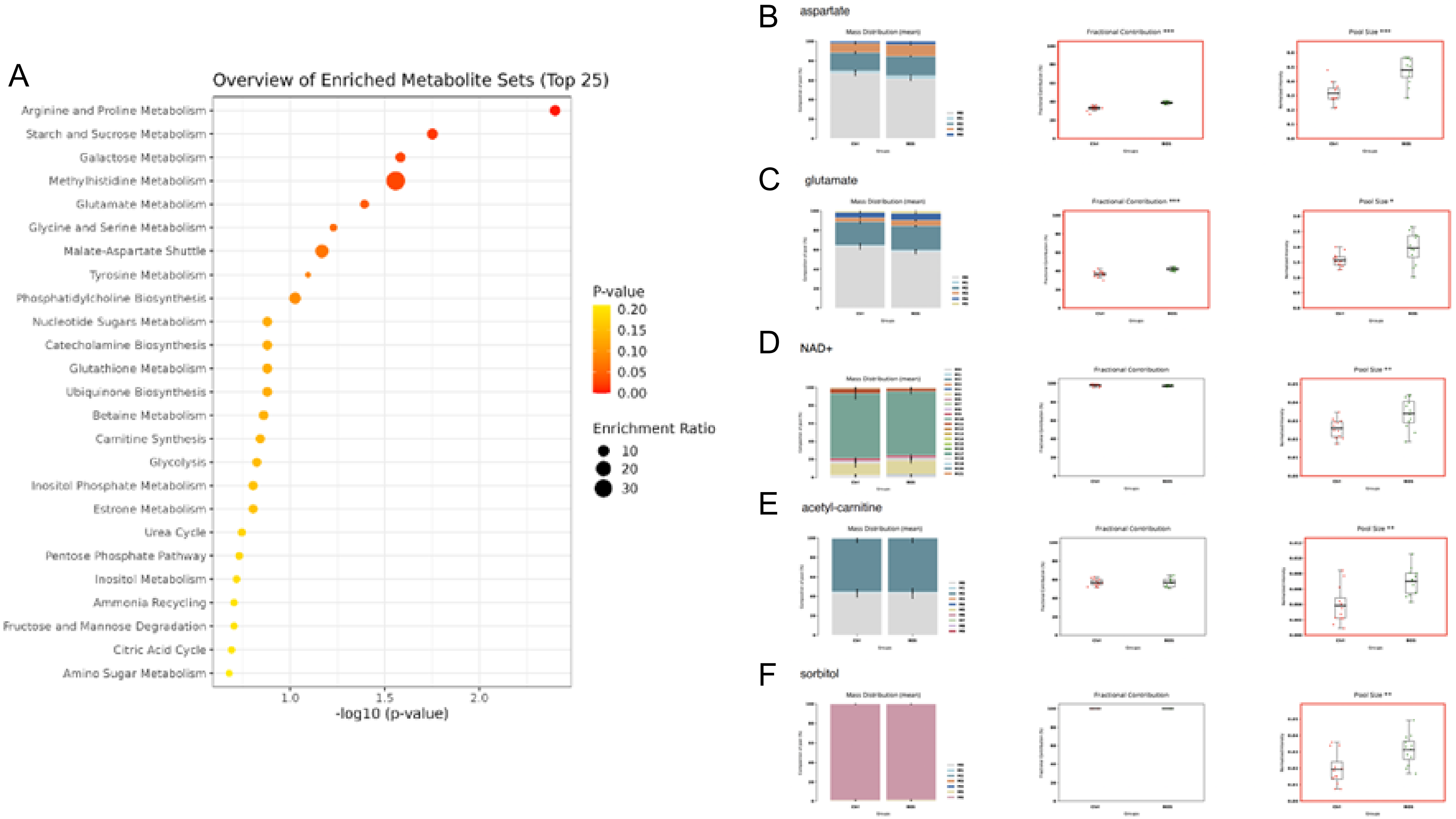

Supp Fig 12: ASXL1 truncation reduces MPC1 protein abundance in HEK293T cells

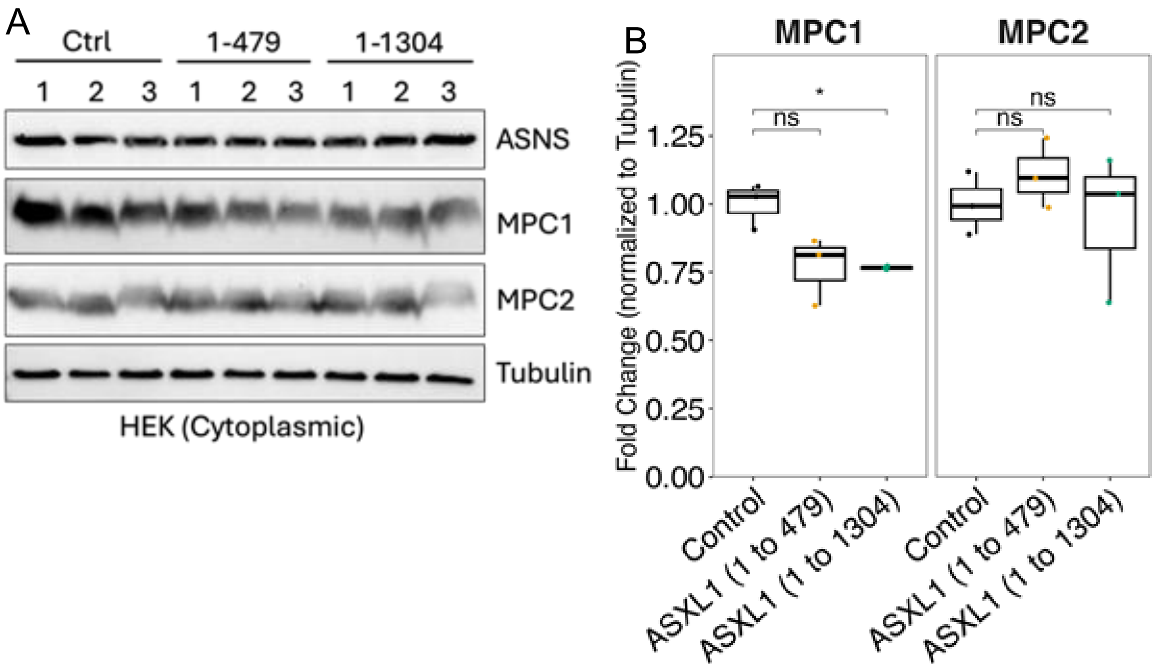

Supp Fig 13: Validation of ASXL1 construct expression in CACO2 cells

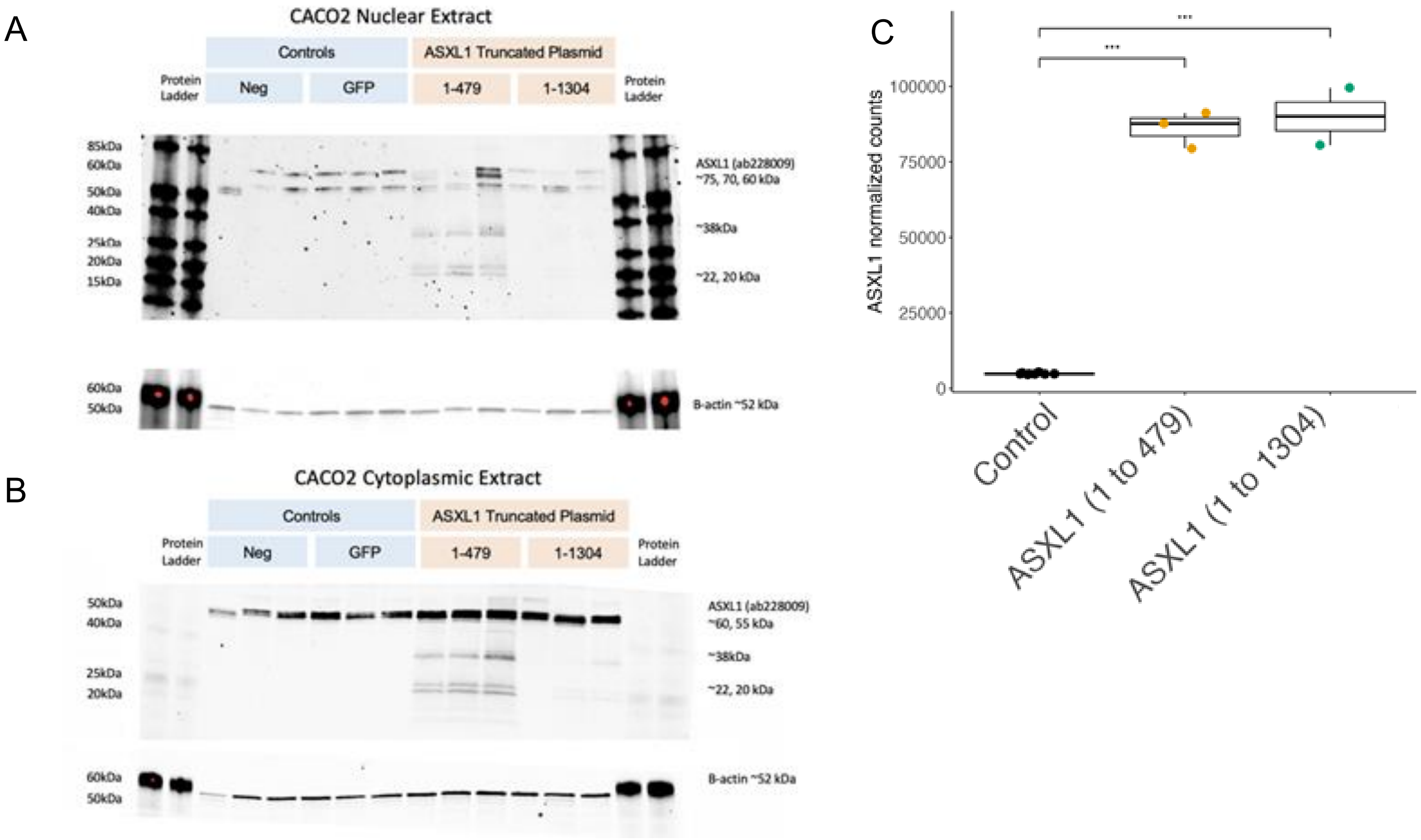

Supp Fig 14: Independent biological replicate experiments confirm metabolic effects of MPC inhibition and Wnt activation

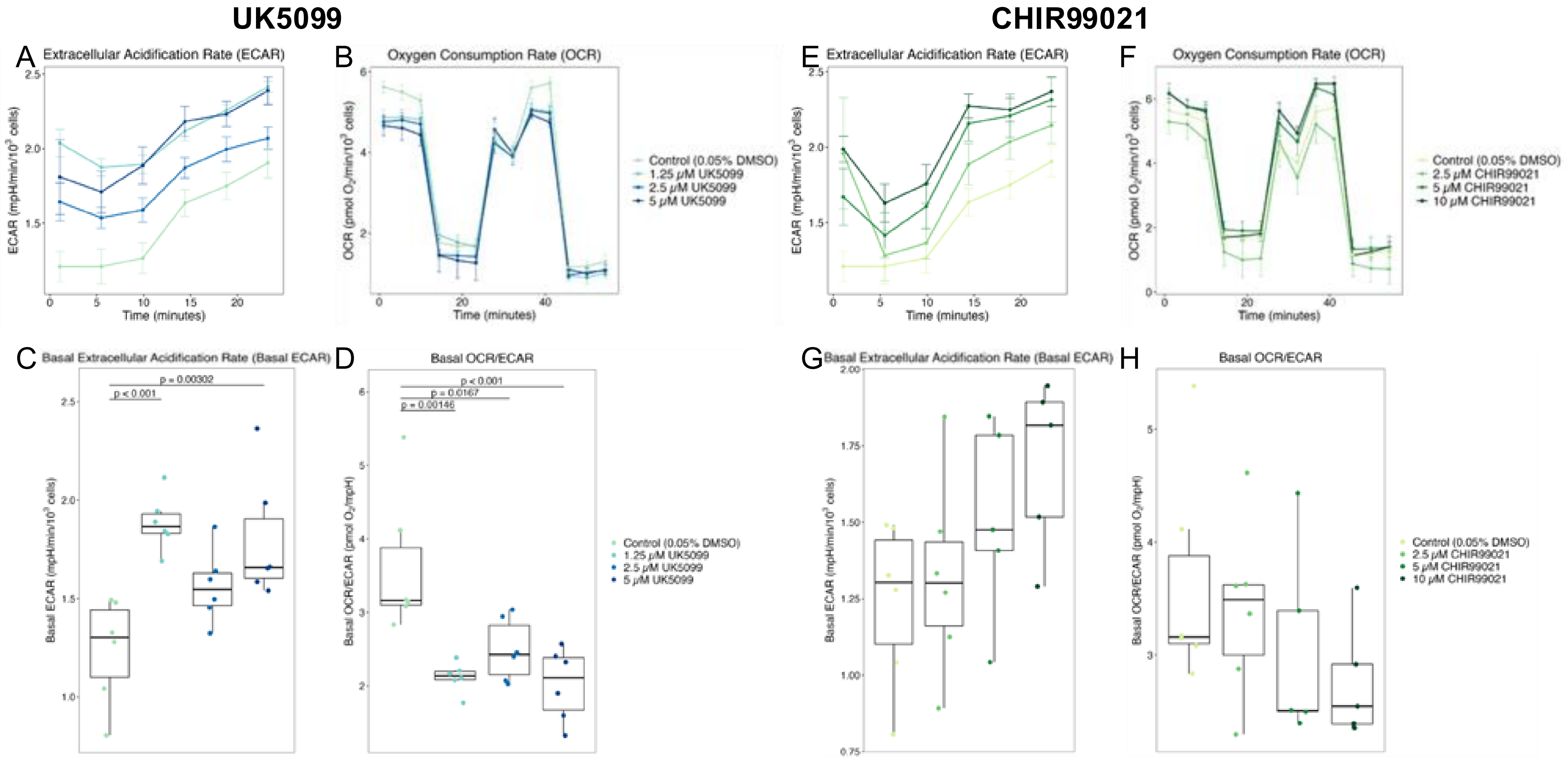

#### Supp Fig 15: BOS fibroblasts exhibit impaired mTOR signaling during amino acid stress

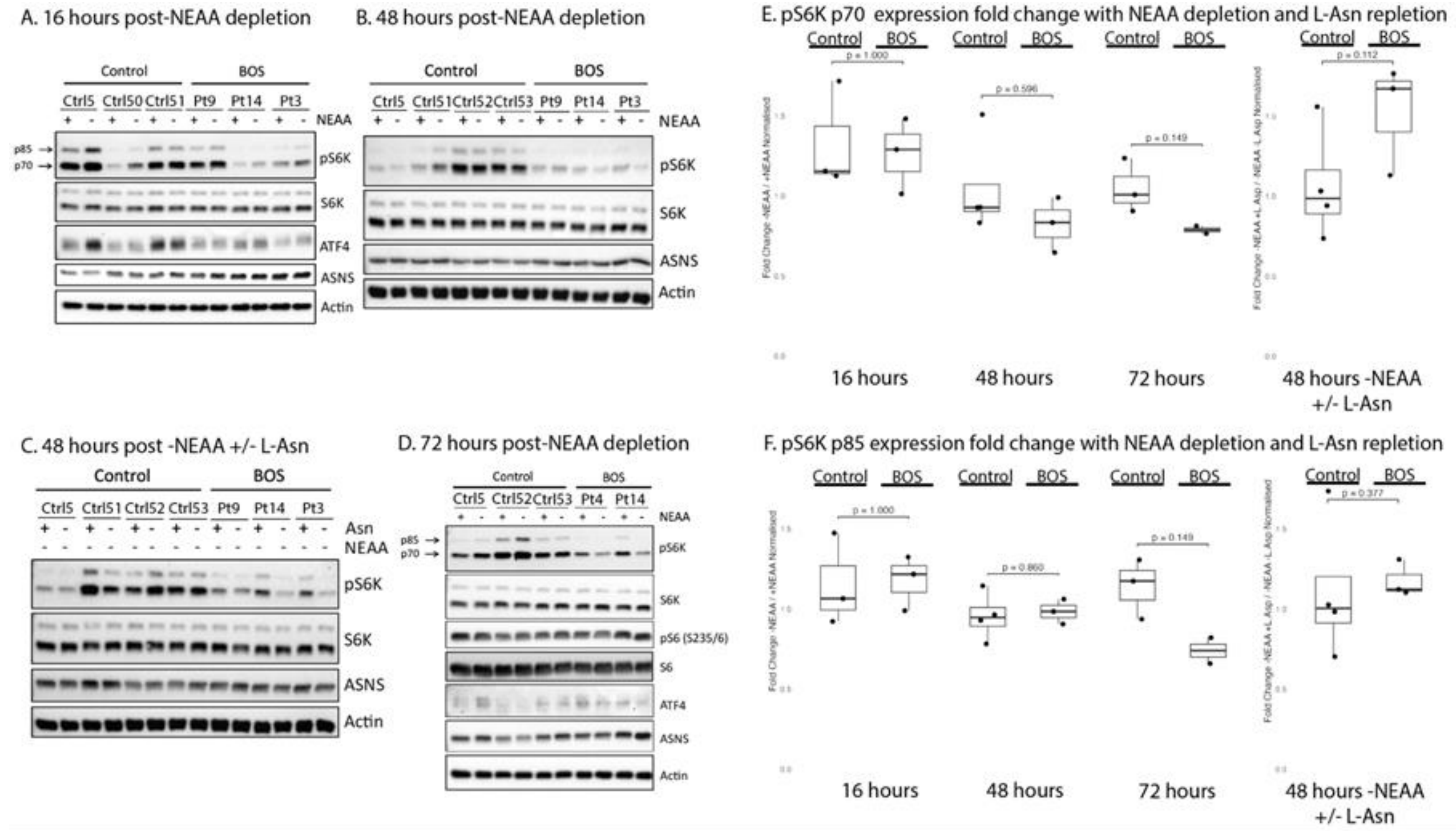

Supp Fig 16: Altered nutrient signaling in BOS fibroblasts occurs despite preserved ATF4 and ASNS transcription

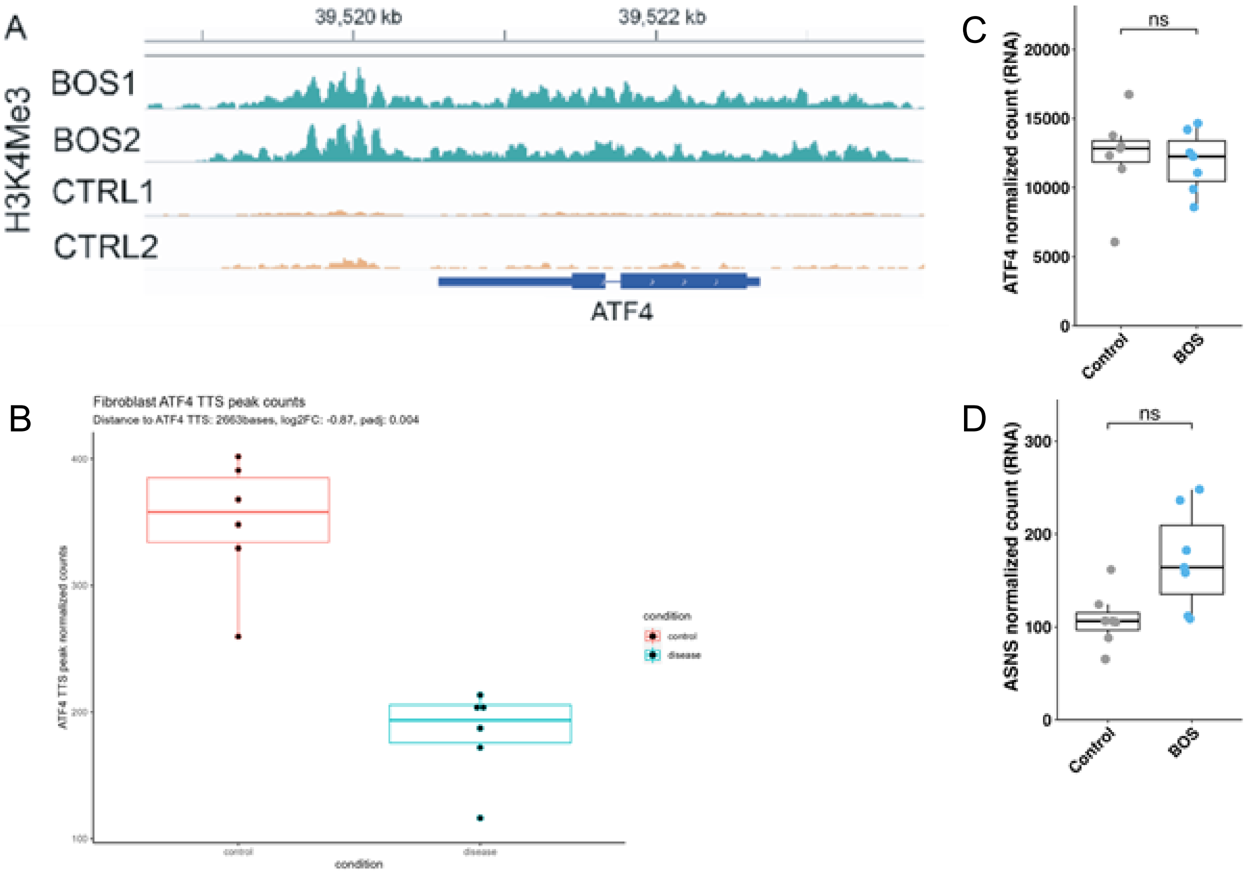

Supp Fig 17: β-catenin does not exhibit differential occupancy at MPC1 and MPC2 promoter regions following Wnt activation

A

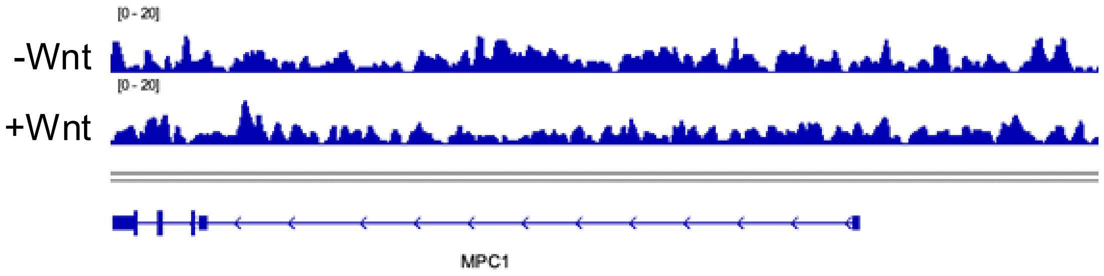


B

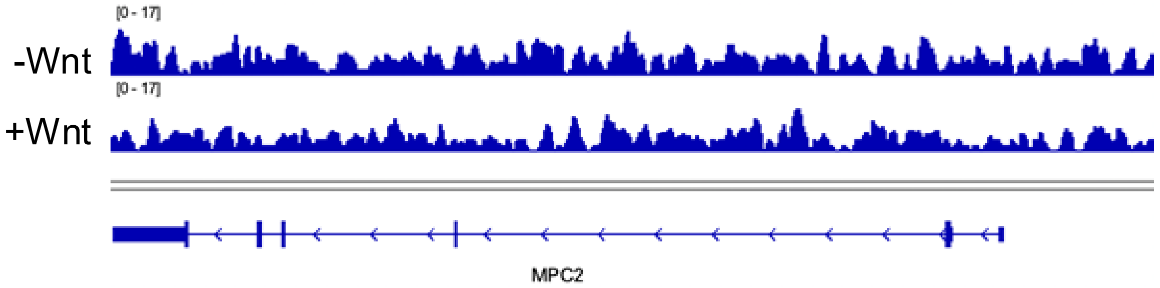


ChIP-seq dataset used: GSE182842

Supp Fig 18: RNA sequencing quality control and transcriptomic validation in HEK293T ASXL1 models

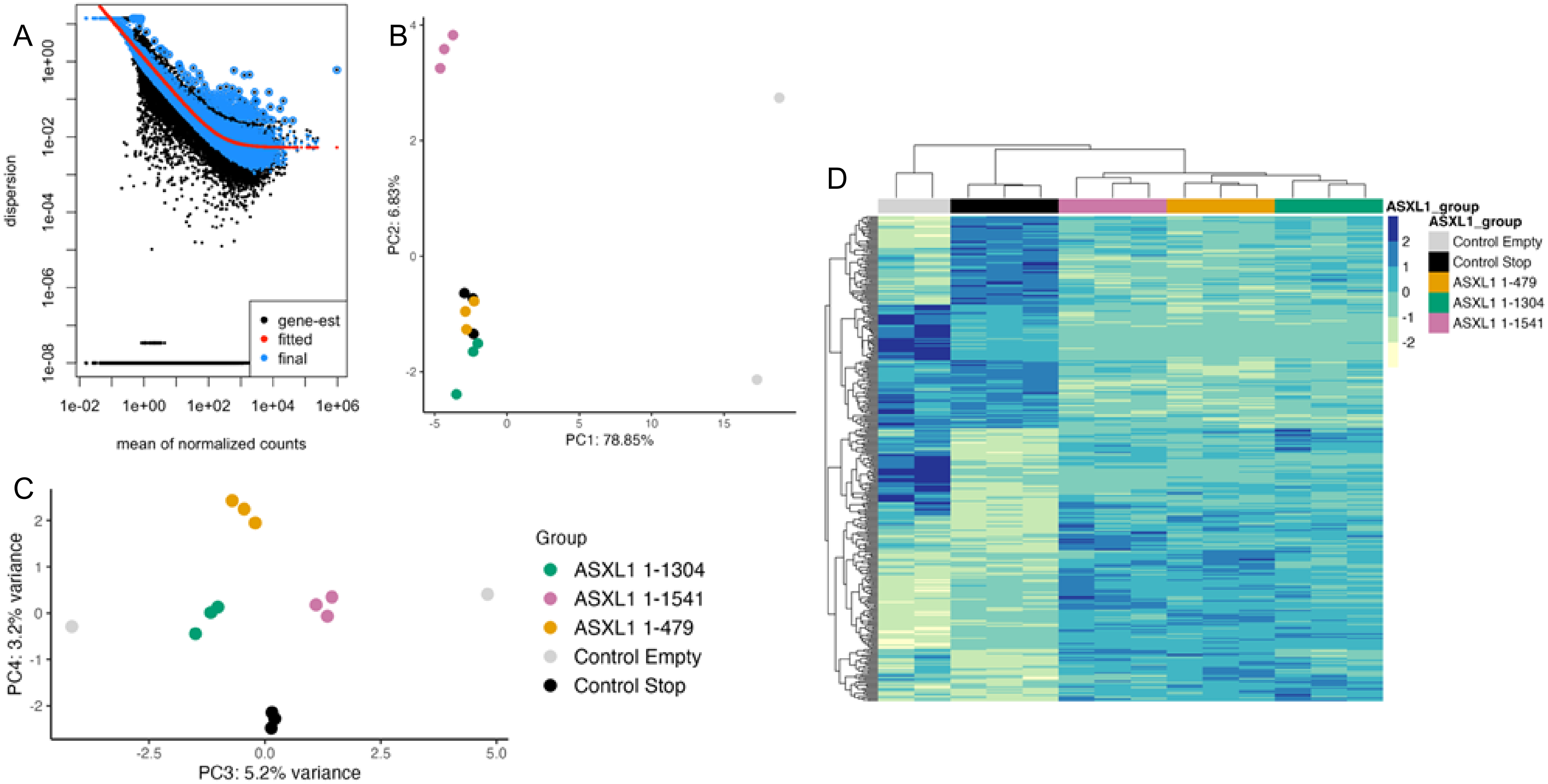

Supp Fig 19: ASXL1 constructs induce distinct transcriptional programs in HEK293T cells

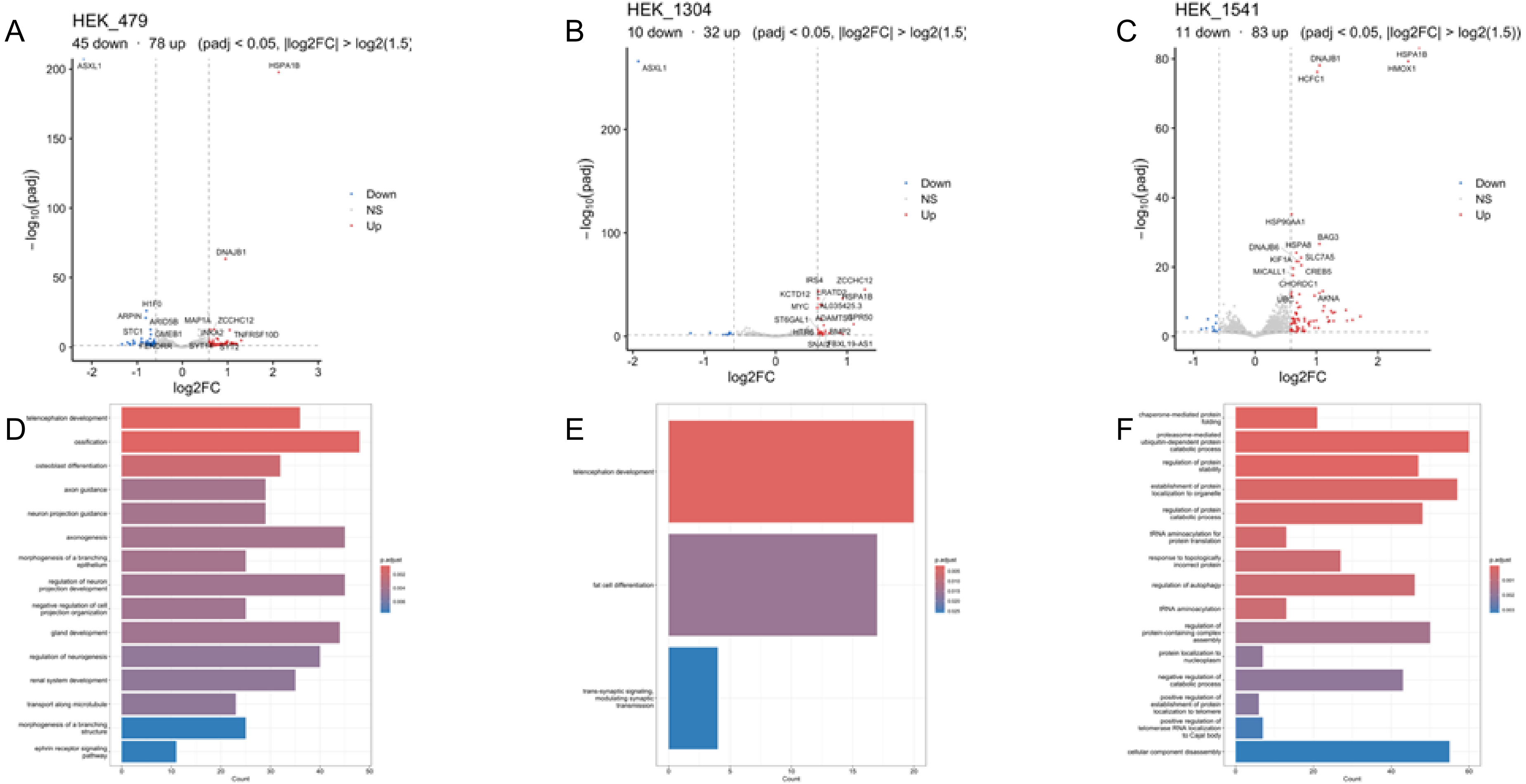

Supp Fig 20: RNA sequencing quality control and transcriptomic validation in CACO2 ASXL1 models

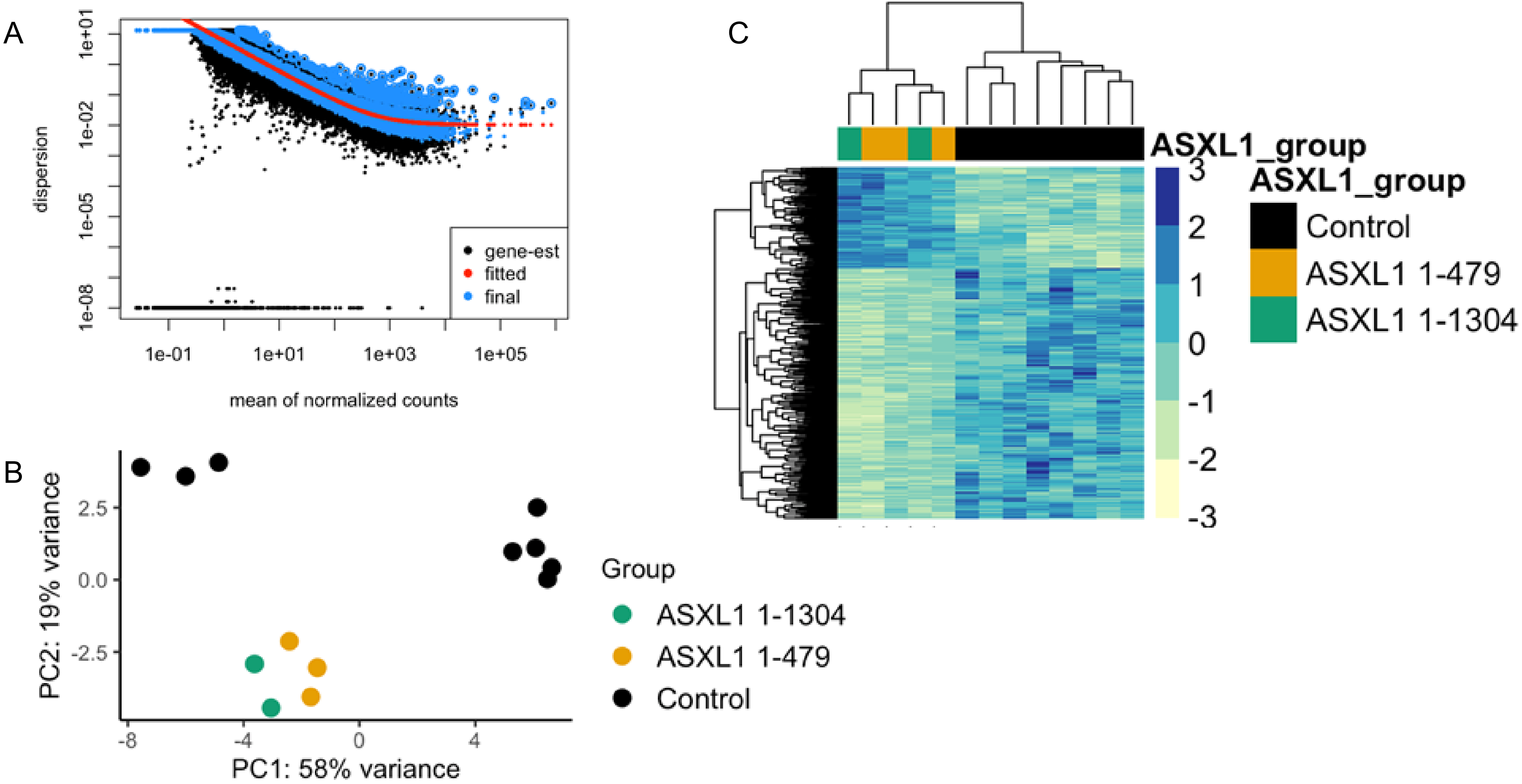

Supp Fig 21: Truncating ASXL1 variants alter mitochondrial and respiratory transcriptional programs in CACO2 cells

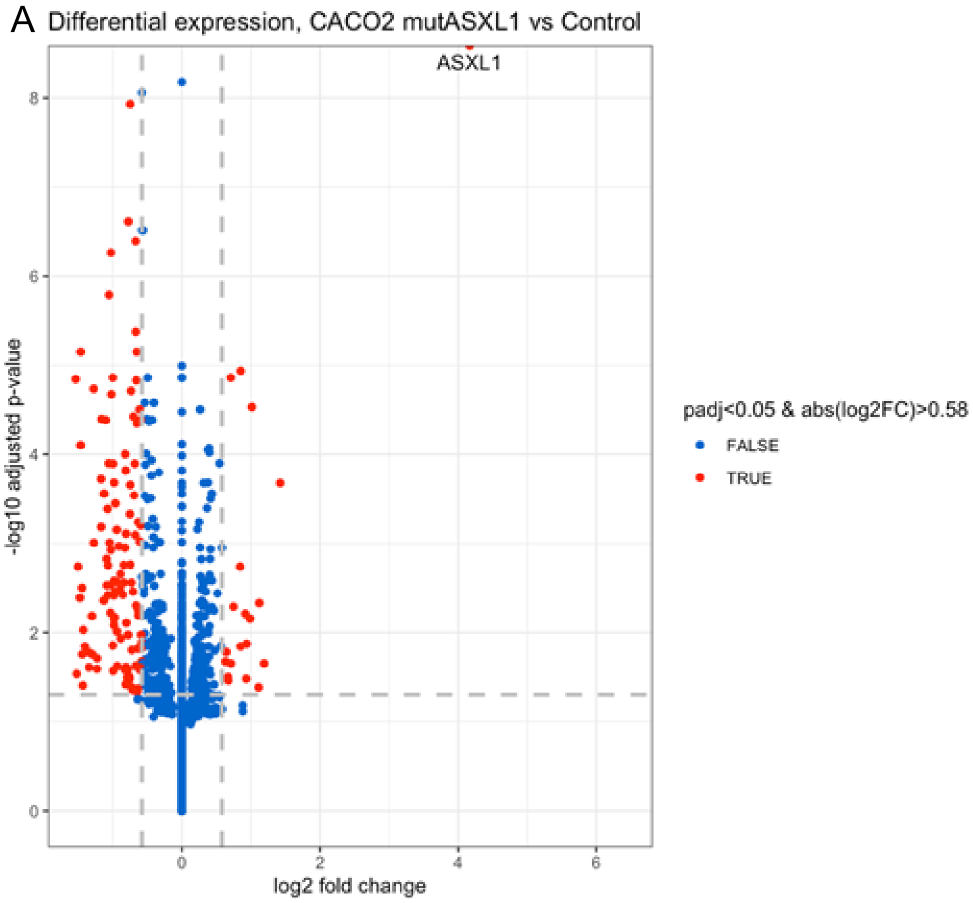


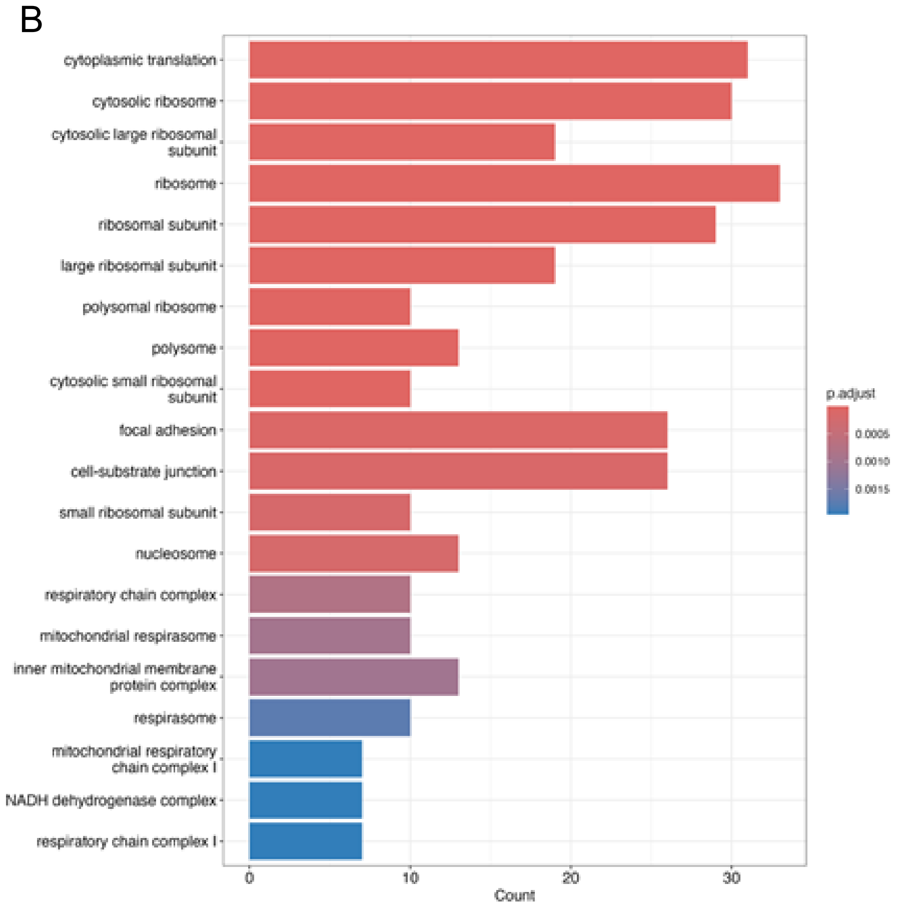

Supp Fig 22: Integrated transcriptomic analysis reveals shared ASXL1-dependent transcriptional signatures across cell types

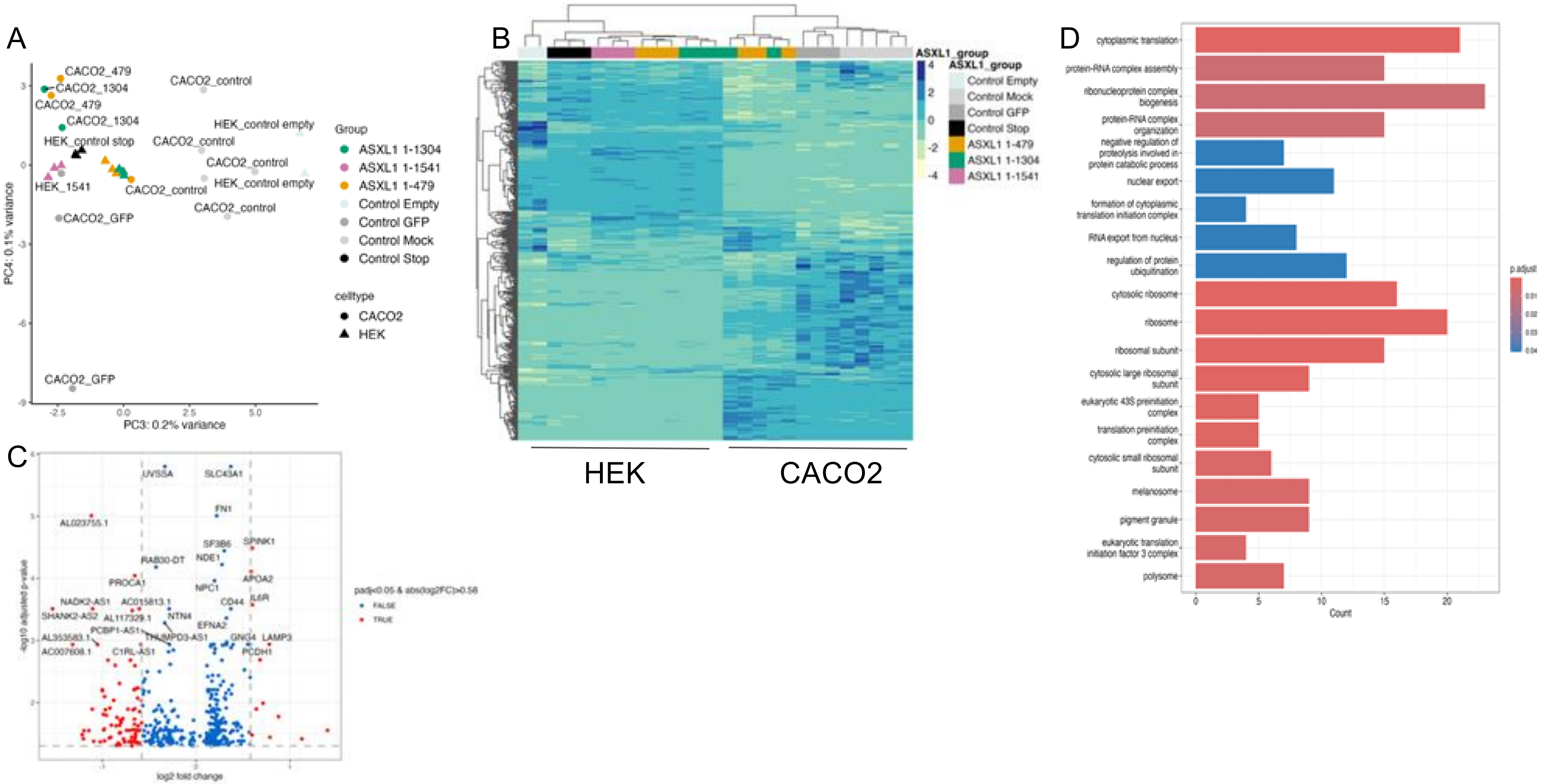

Supp Fig 23: Pyruvate dehydrogenase kinase family genes are not transcriptionally dysregulated following ASXL1 perturbation

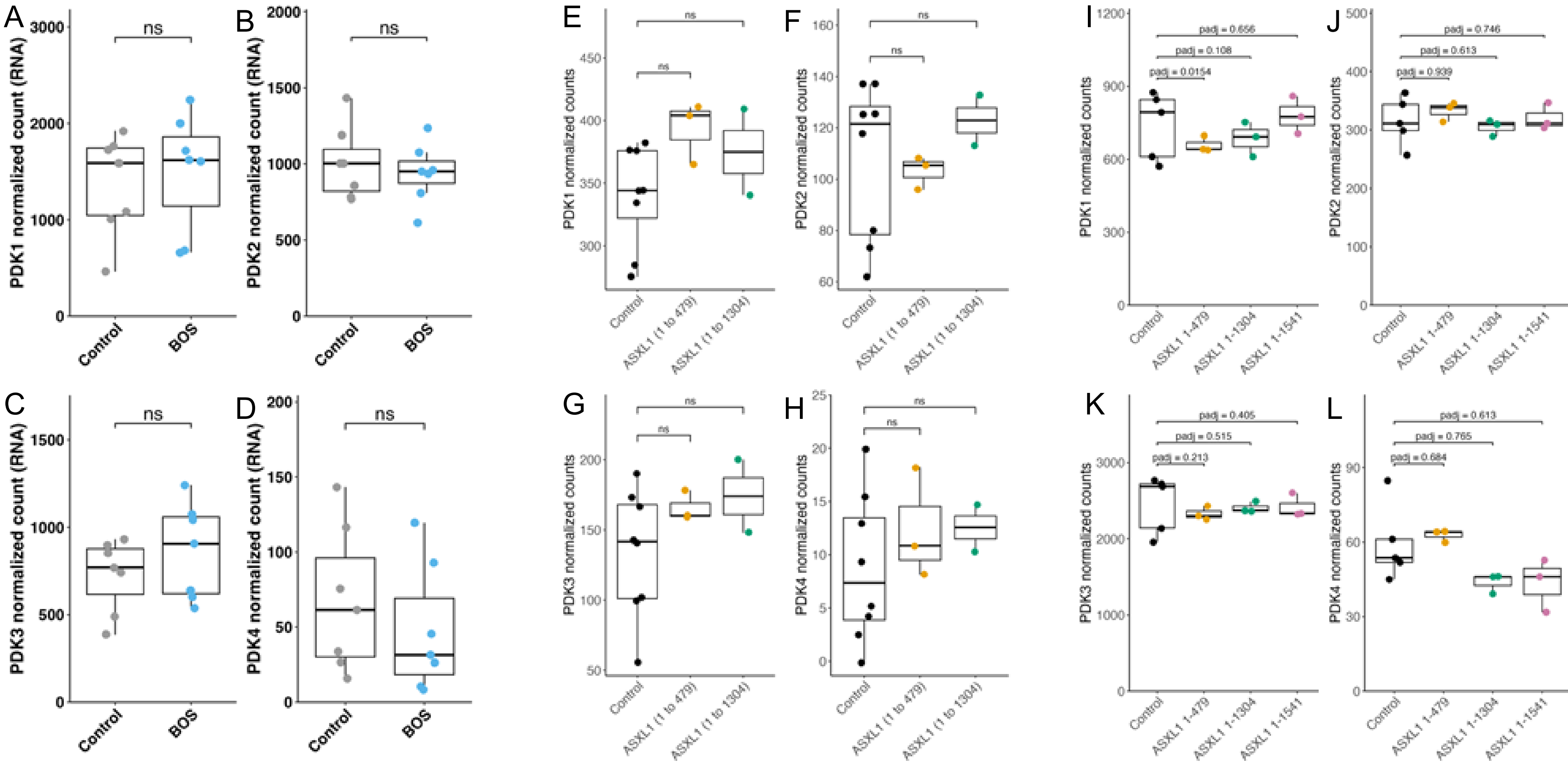

### Supp Fig 24: Glycolytic pathway genes are not transcriptionally dysregulated following ASXL1 perturbation

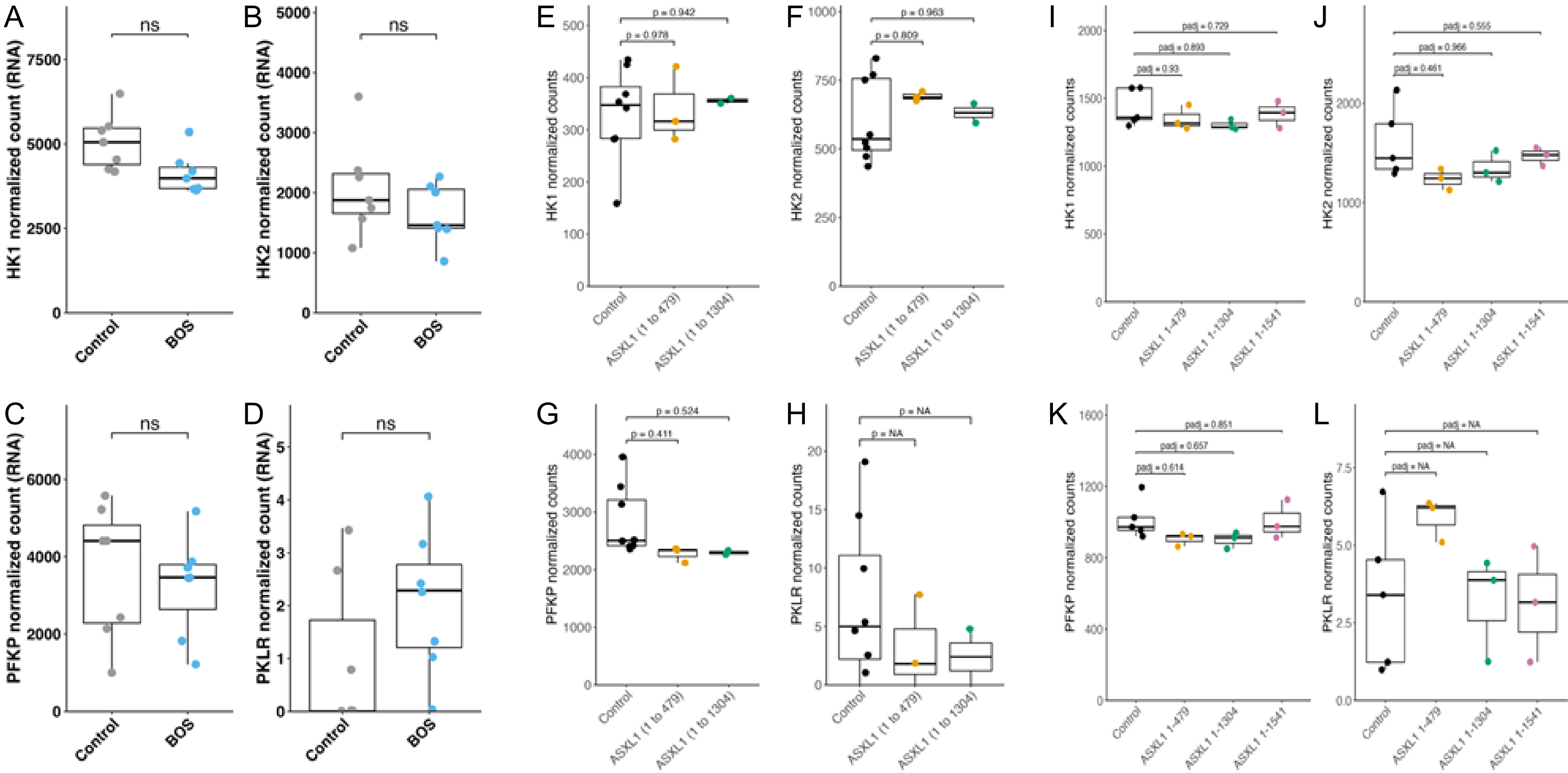

Supp Fig 25: Chromatin accessibility at MPC1 and MPC2 loci is preserved in BOS fibroblasts

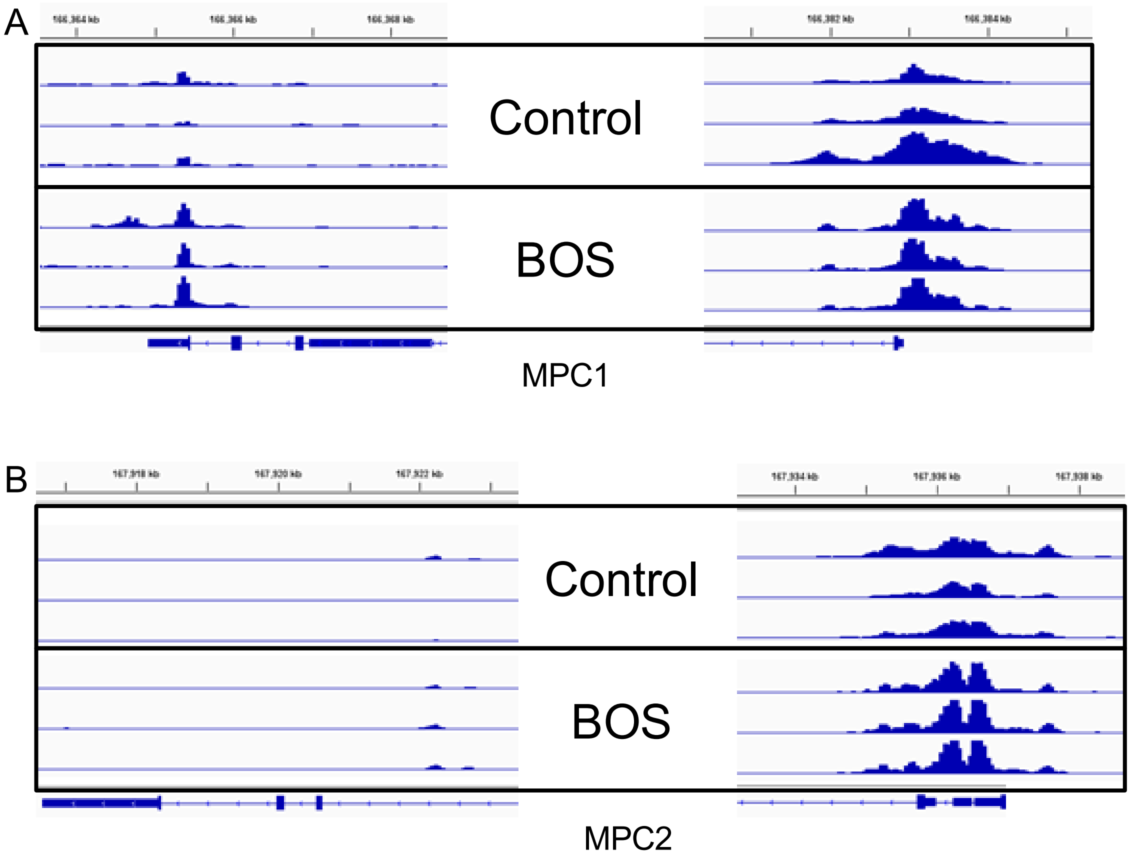
