## Supplemental Table Legends for "Truncating *ASXL1* variants rewire cellular metabolism via mitochondrial pyruvate carrier repression"

**Supplemental Table 1. Seahorse extracellular flux data summary**

Summary of Seahorse extracellular flux experiments from BOS and control fibroblasts, and HEK293T cells transfected with ASXL1 truncating constructs or control plasmids. See **Extended Data** for complete extracellular flux measurements.

**Supplemental Table 2. Metabolomic profiling data summary**

Summary of untargeted and targeted metabolomics, and isotope tracing metabolite experiments. See **Extended Data** for complete normalized data after data filtering.

**Supplemental Table 3. Mummichog pathway enrichment analysis of metabolomic datasets**

Pathway enrichment results generated using Mummichog analysis of metabolomic datasets, including enriched pathways, pathway size, overlap, enrichment score, and statistical significance. Most significant 25 pathways are shown.

**Supplemental Table 4. Metabolomic pathway ontology enrichment analysis**

Gene ontology and pathway enrichment analyses derived from metabolomic datasets identifying biological processes and metabolic pathways altered by ASXL1 truncating variants.

**Supplemental Table 5. Quantification of mitochondrial pyruvate carrier (MPC) protein abundance**

Quantification of MPC1 and MPC2 protein abundance from western blot analyses, including normalization and statistical comparisons across experimental conditions.

**Supplemental Table 6. Quantification of Wnt signaling pathway proteins**

Quantification of proteins associated with Wnt pathway activation, including normalization and statistical analyses across experimental conditions.

**Supplemental Table 7. Quantification of mTOR signaling pathway proteins**

Quantification of proteins associated with mTOR pathway signaling, including normalization and statistical analyses across experimental conditions.

**Supplemental Table 8. Differential gene expression in CACO2 cells expressing ASXL1 plasmid 1–479**

Differential gene expression analysis comparing CACO2 cells expressing ASXL1 plasmid 1–479 and control cells, including log2 fold change, adjusted p value, for the most significantly dysregulated protein-coding genes.

**Supplemental Table 9. Differential gene expression in CACO2 cells expressing ASXL1 plasmid 1–1304**

Differential gene expression analysis comparing CACO2 cells expressing ASXL1 plasmid 1–1304 and control cells, including log2 fold change, adjusted p value, for the most significantly dysregulated protein-coding genes.

**Supplemental Table 10. Gene ontology analysis of transcriptional changes in CACO2 ASXL1 models**

Gene ontology and pathway enrichment analyses of differentially expressed genes identified in CACO2 cells expressing ASXL1 truncating constructs.

**Supplemental Table 11. Differential gene expression in HEK293T cells expressing ASXL1 plasmid 1–479**

Differential gene expression analysis comparing HEK293T cells expressing ASXL1 plasmid 1–479 and control cells.

**Supplemental Table 12. Differential gene expression in HEK293T cells expressing ASXL1 plasmid 1–1304**

Differential gene expression analysis comparing HEK293T cells expressing ASXL1 plasmid 1–1304 and control cells.

**Supplemental Table 13. Differential gene expression in HEK293T cells expressing ASXL1 plasmid 1–1541**

Differential gene expression analysis comparing HEK293T cells expressing ASXL1 plasmid 1–1541 and control cells.

**Supplemental Table 14. Gene ontology analysis of transcriptional changes in HEK293T ASXL1 models**

Gene ontology and pathway enrichment analyses of differentially expressed genes identified in HEK293T cells expressing ASXL1 truncating constructs.

**Supplemental Table 15. Shared transcriptional alterations across CACO2 and HEK293T ASXL1 models**

Differentially expressed genes shared between CACO2 and HEK293T ASXL1 models.

**Supplemental Table 16. Conserved gene ontology signatures across ASXL1 cellular models**

Gene ontology and pathway enrichment analyses identifying conserved biological pathways altered across CACO2 and HEK293T ASXL1 models.

**Supplemental Table 17. Catalogue of reagents and materials**

List of reagents, chemicals, kits, and experimental materials used in this study, including vendor information and catalogue numbers.

**Supplemental Table 18. Antibodies used for immunoblotting analyses**

Primary and secondary antibodies used in western blot analyses.

**Supplemental Table 19. Primers used for ASXL1 plasmid construction and validation**

Primer sequences used for ASXL1 construct generation, cloning, and validation.
